# Dynamic action of the Sec machinery during initiation, protein translocation and termination revealed by single molecule fluorescence

**DOI:** 10.1101/248310

**Authors:** Tomas Fessl, Daniel Watkins, Peter Oatley, William J. Allen, Robin A. Corey, Jim E. Horne, Steve A. Baldwin, Sheena E. Radford, Ian Collinson, Roman Tuma

**Affiliations:** Astbury Centre for Structural Molecular Biology, University of Leeds; School of Molecular and Cellular Biology, Faculty of Biological Sciences, University of Leeds, LS2 9JT, UK; School of Biochemistry, University of Bristol, BS8 1TD UK; School of Biomedical Sciences, Faculty of Biological Sciences, University of Leeds, LS2 9JT, UK; Faculty of Science, University of South Bohemia, Ceske Budejovice, Czech Republic

**Author notes:** Current Address: Department of Molecular Biology and Biotechnology, University of Sheffield, S10 2TN, UK.

## Abstract

Protein translocation across cell membranes is a ubiquitous process required for protein secretion and membrane protein insertion. This is mediated, for the majority of proteins, by the highly conserved Sec machinery. The bacterial translocon – SecY_MK_EG – resides in the plasma membrane, where translocation is driven through rounds of ATP hydrolysis by the cytoplasmic SecA ATPase, and the proton motive force (PMF). We have used single molecule Förster resonance energy transfer (FRET) alongside a combination of confocal and total internal reflection microscopy to gain access to SecY pore dynamics and translocation kinetics on timescales spanning milliseconds to minutes. This allows us to dissect and characterise the translocation process in unprecedented detail. We show that SecA, signal sequence, pre-protein and ATP hydrolysis each have important and specific roles in unlocking and opening the Sec channel, priming it for transport. After channel opening, translocation proceeds in two phases: an initiation phase independent of substrate length, and a length-dependent transport phase with an intrinsic translocation rate of ~ 40 amino acids per second for the model pre-protein substrate proOmpA. The initiation and translocation phases are both coupled to ATP hydrolysis while termination is ATP-independent. Distributions of translocation rates reflect the stochastic nature of the translocation process and are consistent with the recently proposed Brownian ratchet model [Allen *et al.* doi: 10.7554/eLife.15598]. The results allow us unparalleled access to the kinetics of the complex reaction and provide a framework for understanding the molecular mechanism of protein secretion.

## Introduction

Protein secretion is an essential and conserved process in all living cells, responsible for the delivery of proteins to the cell surface. The major route for this process is *via* the ubiquitous Sec machinery (Collinson et al., 2015), which comprises at its core a heterotrimeric complex: SecYEG in the plasma membrane of bacteria and archaea, and Sec61αβγ in the eukaryotic endoplasmic reticulum (ER). Pre-proteins are targeted to the Sec machinery by an N-terminal signal sequence (SS) or a transmembrane helix (TMH), and translocated through the Sec machinery in an unfolded conformation. This can occur either during their synthesis (co-translationally), or afterwards (post-translationally); in the latter case, pre-proteins are maintained in an unfolded state in the cytoplasm prior to secretion by chaperones such as SecB (Kumamoto and Beckwith, 1983; Weiss et al., 1988). In bacteria, inner membrane proteins are generally secreted co-translationally, while proteins destined for the periplasm, outer membrane or extracellularly, tend to follow the post-translational route, driven by the ATPase SecA (Hartl et al., 1990); (Brundage et al., 1990).

The protein channel is formed through the centre of SecY, between two pseudo-symmetrical halves, each containing five TMHs (Cannon et al., 2005; Van den Berg et al., 2004) (Fig. 1). Separation of these domains opens a channel across the membrane (secretion) as well as a lateral gate (LG) for SS docking and membrane protein departure (insertion) into the membrane (Fig. 1B). When at rest the channel is kept closed by a short, usually α-helical plug and a ring of six hydrophobic residues (Van den Berg et al., 2004) to prevent ion leakage and dissipation of the PMF (Saparov et al., 2007). Activation is achieved by the ribosome nascent chain complex (Jomaa et al., 2016; Voorhees and Hegde, 2016), or by the association of the pre-protein, SS, and the translocating ATPase SecA in bacteria (Lill et al., 1989) or BiP in the ER lumen of eukaryotes (Nguyen et al., 1991).

**Figure 1:**
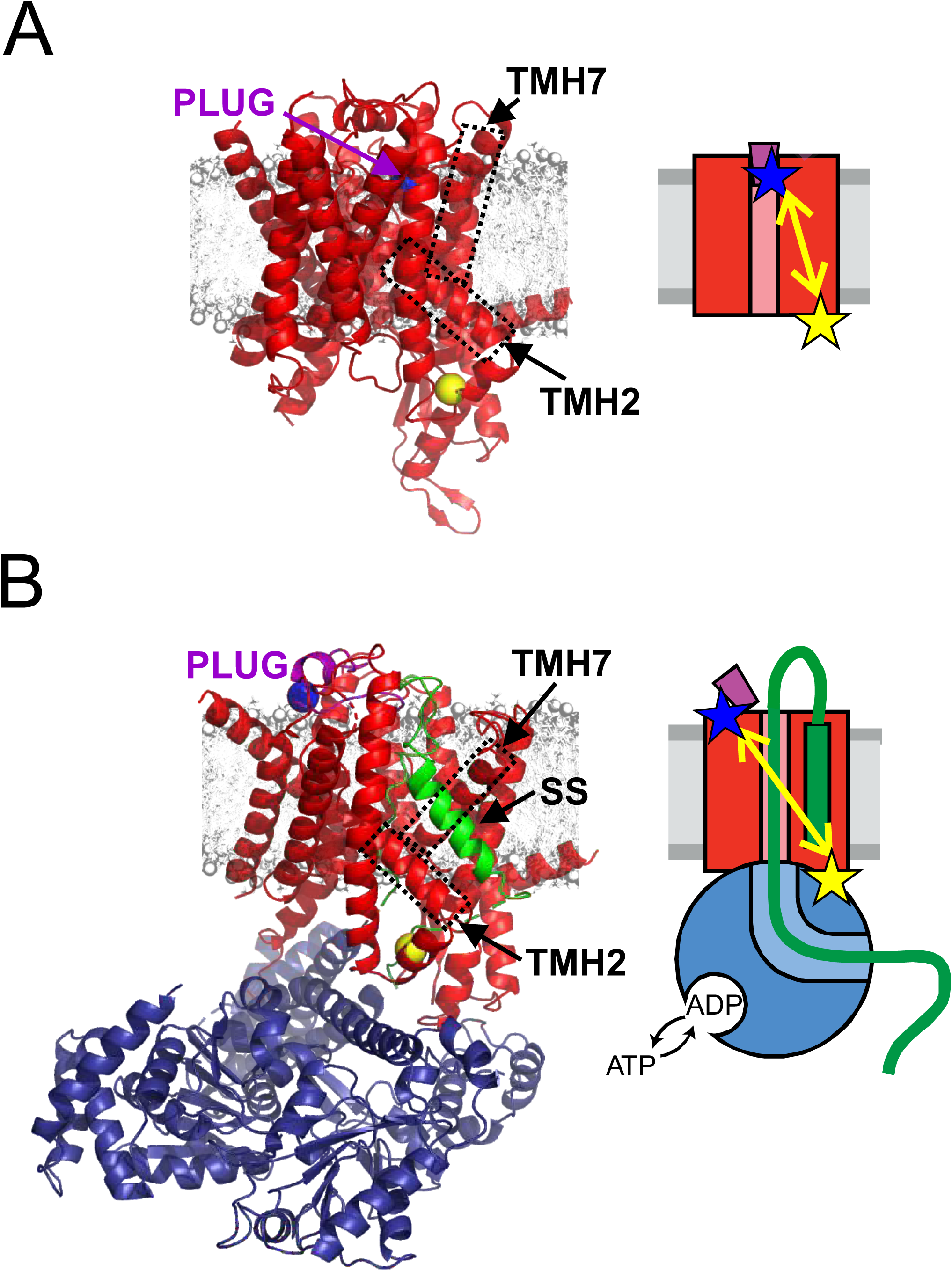
Channel opening and helical plug motion illustrated by available high-resolution structures. **A:** Closed SecYEG (PDB: 5AWW, (Tanaka et al., 2015)). SecYEG (red), modelled membrane (grey), plug helix (purple, signal sequence (SS) is green, transmembrane). Transmembrane helices TMH2 and TMH7 are identified by dashed boxes. The residue M63 within the plug is depicted as blue ball while a cytoplasmic side reference residue K106 is shown as a yellow ball. A schematic of each state is shown on the right with SecYEG in red and SecA in blue, the plug in purple and translocated polypeptide green. The respective distances are shown as yellow double-arrow line with the dyes shown as stars (blue and yellow). **B:** The open state SecYEG:SecA (PDB: 5EUL, (Li et al., 2016). Colours and labelling are as in panel A, with Sec A (Blue) and the signal sequence (SS) in green.

Despite numerous protein structures of the bacterial, archaeal and eukaryotic Sec systems, (see (Collinson et al., 2015) and references therein), no consensus has yet emerged as to the dynamic mechanism underlying protein translocation through the Sec protein pore. Post-translational translocation in bacteria is initiated by binding of SecA and pre-protein to SecYEG, and insertion of the SS into the LG of SecY – at the interface with the lipid bilayer, between TMHs 2 and 7 (Fig. 1B) (Briggs et al., 1986; Hizlan et al., 2012; Li et al., 2016; McKnight et al., 1991). Several conformational changes accompany this insertion: TMH7 is relocated to make room for the SS, and the plug is displaced from the channel, in a process termed ‘unlocking’ (Corey et al., 2016b; Hizlan et al., 2012). Meanwhile, binding of SecA causes a partial opening of the protein channel in SecY and the mobilisation of the protein cross-linking domain of SecA, which forms a clamp around the translocating pre-protein (Zimmer et al., 2008). Together, these steps lead to full activation of the SecA ATPase and prime the channel to translocate the remainder of the pre-protein. However, the order of events, energy requirements and kinetics of these steps have yet to be convoluted.

There is less agreement in the field as to the mechanism of translocation itself. Two principal models have been proposed: (1) a processive power-stroke in which a fixed length of substrate is transported for each ATP molecule hydrolysed by SecA (Erlandson et al., 2008; Zimmer et al., 2008); and (2) a Brownian ratchet model in which passive diffusive motion is conferred directionality by gating at the expense of ATP hydrolysis (Allen et al., 2016; Liang et al., 2009). Recent studies favour at least some element of diffusion: previously, we proposed a “pure” stochastic model, in which the free energy available from ATP binding and hydrolysis at SecA drives a Brownian ratchet at the SecY LG (Allen et al., 2016), while others have suggested a hybrid processive/stochastic model, in which ATP hydrolysis generates a power stroke (‘push’) on the polypeptide, but the latter is allowed to diffuse through the pore (‘slide’) (Bauer et al., 2014). This marks a shift from deterministic models based on static structural snapshots to a stochastic view in which intrinsic dynamics of the complex are taken into account (Corey et al., 2016a).

A central problem in addressing these questions is the challenge of dissecting the rates of the various steps, and their dependence on extrinsic factors such as ATP concentration or pre-protein sequence. For example, quantification of the protein translocation rate has been undermined by problems inherent to ensemble analysis of unsynchronised reactions; wherein multiple steps are convoluted into one overall measureable value; *i.e.* an overall translocation rate or average rate of ATP hydrolysis (Brundage et al., 1990; De Keyzer et al., 2002). Furthermore, many of these ‘translocation rates’ are determined from sparsely populated, discontinuous assays not quenched instantly, but stopped by the addition of protease. This situation may explain why wildly different figures have been published for the energetic cost of transport: one study proposed a single ATP to drive the passage of ~ 5 kDa of protein (roughly 40 amino acids) across the membrane (van der Wolk et al., 1997); while a later analysis (from the same lab) arrived at 5 molecules of ATP hydrolysed per single amino acid transported (Tomkiewicz et al., 2006).

Here, we exploit single molecule Förster resonance energy transfer (FRET) analyses – building on a previous study (Allen et al., 2016) – to dissect the mechanism of protein translocation in unprecedented detail. This approach, which utilises an array of single molecule FRET experiments sensitive to different timescales alongside ensemble measurements, allowed us to delineate several stages of translocation: (1) SS-dependent but ATP-independent unlocking of the translocon; (2) ATP-dependent plug opening; (3) a pre-processive translocation stage; (4) ATP-dependent processive translocation and (5) ATP-independent, fast channel closing. This has enabled us to estimate an intrinsic, processive translocation rate of ~40 amino acids per second. The broad distribution of translocation rates is consistent with the stochastic Brownian ratchet model (Allen et al., 2016).

## Results

### Selection of surface residues for dye attachment

Building on the successful application of single molecule FRET to follow SecYEG LG opening (Allen et al., 2016), we utilised a similar approach to follow another key event associated with protein transport: the movement of the SecY plug during complex activation and protein channel opening. The plug helix is expected to relocate during activation of the channel by association of the SS and SecA, and remains open during the protein translocation process (Li et al., 2016; Zimmer et al., 2008);(Bieker et al., 1990; Flower et al., 1995; Hizlan et al., 2012; Robson et al., 2009a; Tam et al., 2005). Met63 (*E. coli* numbering) of the plug region of SecY was selected for dye attachment in order to monitor its mobility (Figure 1). As FRET is most sensitive for inter-dye distances close to the Förster radius (6 nm for Alexa Fluor 488 and 594 dye pair used here), we chose the solvent-accessible residue Lys106, within the loop on the cytoplasmic side of SecY, as the reference dye attachment site (Figure 1).

In order to verify the suitability of this labelling scheme, the positions accessible to the attached dyes (accessible volumes) were modelled onto the available closed and open state crystal structures using a trial and error (Monte Carlo) protocol that checks for steric clashes (Figure 1S1A-D). The resulting inter-dye distance distributions (Figure 1SE) yielded theoretical FRET efficiency (E_FRET_) histograms (Figure 1S1F) centred at low E_FRET_ values (0.16-0.2) for the two open configurations sampled here and at 0.4 with ~0.6 shoulder for the closed state. These differences are sufficient to be distinguishable by single molecule FRET; hence, M63 and K106 were mutated to cysteine in a Cys-free SecYEG variant (Deville et al., 2011) and the resulting protein was labelled with Alexa Fluor 488 and 594 maleimide dyes (the doubly labelled protein is hereafter designated SecY_MK_EG). Bulk protease protection transport assays with SecY_MK_EG reconstituted into proteoliposomes indicated that the labelled variant remained active and translocated a model preprotein substrate, proOmpA delta 176 (pOA 176), into the lumen of the vesicles with an efficiency akin to that of WT (Figure 1S2).

### Single molecule monitoring of plug opening

The expected E_FRET_ signal during translocation is schematically depicted in Figure 2A. The duration (dwell time) of the open, low-FRET state is related to the duration of translocation event and thus is expected to be inversely dependent on the translocation rate and to increase with the length of the substrate. Based on previous ensemble translocation rate estimates (Brundage et al., 1990; De Keyzer et al., 2002), the open state is expected to last from seconds to minutes, while the transitions between the open and closed states (Figure 2A, red and blue dashed vertical lines) although unknown, are likely to be much faster. In order to capture the slow dwell times and potentially fast transitions, two complementary single molecule detection techniques were employed: confocal microscopy for detection of events on the millisecond time scale (Figure 2B) and total internal reflection (TIRF) imaging of immobilised vesicles for much longer observations lasting up to several minutes (Figure 2C). The former allowed us to explore the rate of conversion between closed and open states, whereas the latter was used to measure the duration of translocation and determine the translocation rate for individual SecY_MK_EG complexes.

**Figure 2:**
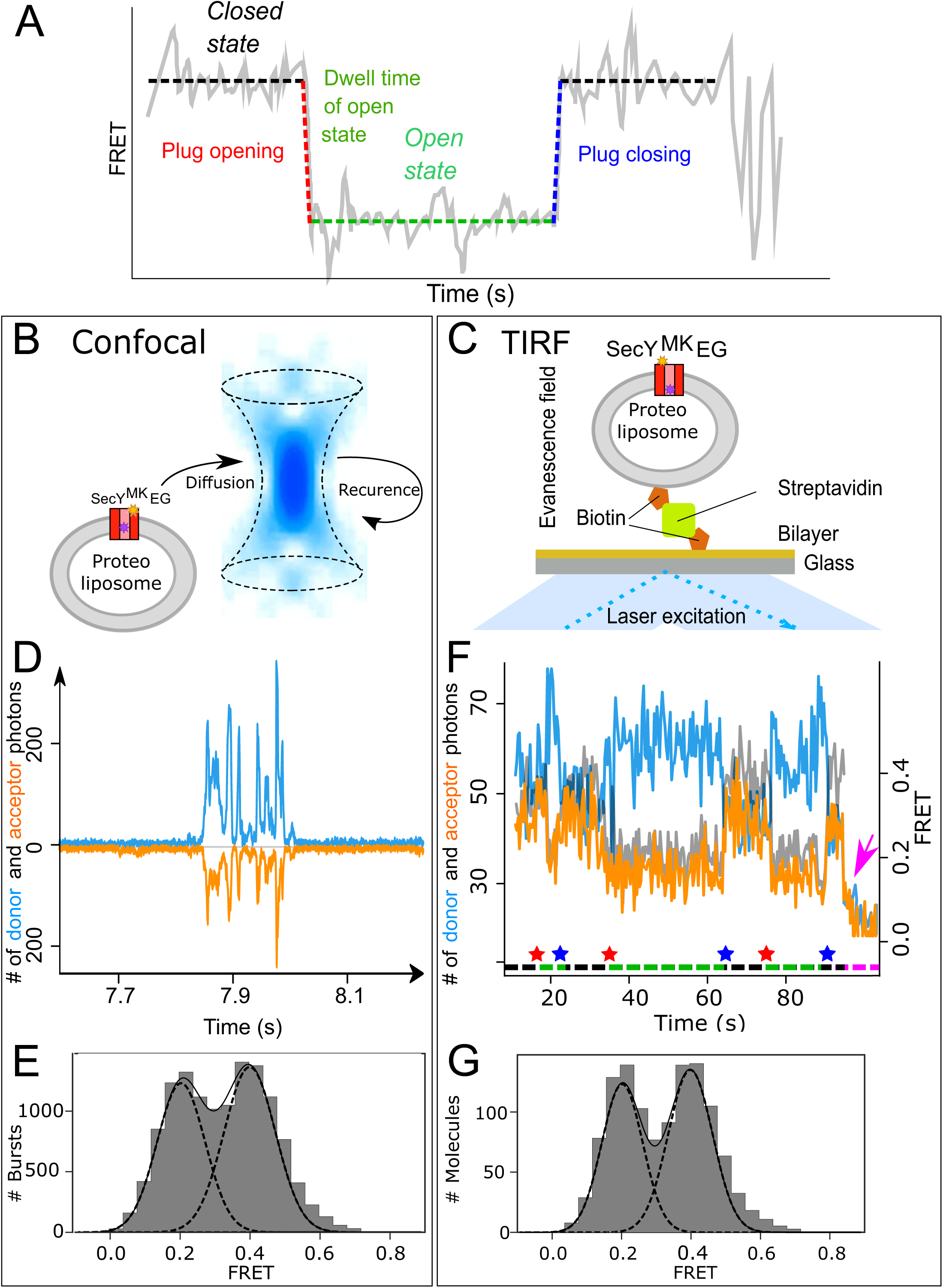
Monitoring plug opening by single molecule FRET. **A:** Expected changes in the FRET efficiency as a consequence of plug opening during translocation. Pre-translocation, high FRET closed state (black dashed line) changes rapidly to a low FRET, open state (red dashed line) and remains open (green dashed line) until closing (blue dashed line). **B:** Schematic depiction of confocal (blue confocal volume) detection of freely diffusing proteoliposomes containing SecY_MK_EG (red) embedded in the bilayer (grey) with a recurrence path (arrow). **C:** Schematic depiction of proteoliposome immobilized via a biotinylated lipid to a streptavidin (green) coated cover slip. Laser beam (blue) in a total internal reflection fluorescence (TIRF) mode creates a thin layer (~500 nm) of evanescent optical field close to the surface (blue gradient). **D:** Example of fluorescence time traces collected in confocal microscope (donor channel is blue, acceptor orange shown in opposite orientation for clarity) containing a train of bursts from recurrence. FRET data sets were collected in the presence of short pOA substrate (100 aa, pOA100, 700 nM), the ATPase SecA (1 µM), the chaperone SecB (10 µM) and 2 mM ATP. **E:** FRET efficiency histograms derived from confocal data (10,000 events). A sum (solid black line) of two Gaussian functions (black dashed lines) approximates the experimental histograms. The histogram was corrected for contribution from the 50% SecY_MK_EG in opposite orientation which is unable to bind SecA and translocate as described in Supplementary Figure 2S1. **F:** Example of TIRF fluorescence trace for translocation of pOA100 with dwell times on the order of seconds (donor channel is blue, acceptor orange and FRET efficiency shown in grey). The system starts in a closed state, undergoes initiation and opening of the plug (indicated by a red star below the trace) which remains open during translocation (green dashed line under the trace). After translocation is finished, the plug snaps back (blue star) to seal the pore and the system remains in the closed state (black dashed line) until another round of translocation or one of the dyes photobleaches (magenta arrow). Note that duration of the translocation events varies and reflects the stochastic nature of the process. **G:** TIRF data histogram (300 events) collected during translocation of pOA100. Fitting to two Gaussians is depicted as in the panel E.

For single molecule analysis, SecY_MK_EG was reconstituted into 100 nm diameter proteoliposomes formed from *E. coli* polar lipids, at concentrations to give at most one molecule of translocon per vesicle (Allen et al., 2016; Deville et al., 2011). The SecY_MK_EG proteoliposomes were then either diluted to pM concentration for confocal detection (Figure 2B) or immobilised onto a glass surface via a biotin:streptavidin linker and imaged using TIRF microscopy (Figure 2C). For each setup, FRET datasets were collected in the presence of all components required for translocation: pOA, the ATPase SecA, the chaperone SecB and ATP. Translocation events are expected to appear as transient reductions in E_FRET_, as the plug moves away from the channel (see Fig. 2A).

With confocal detection, the time-dependent fluorescent intensity traces consist of a series of short signal bursts (Figure 2D) that correspond to the passage of individual vesicles through the confocal volume. The duration of each burst is limited by diffusion to a few milliseconds. Bursts were extracted from the signal traces, converted to FRET efficiencies (see Methods) and collated in a histogram (Figure 2E). The result exhibits bimodal distribution of the expected FRET efficiencies – E_FRET_~0.2 and ~0.4.

The TIRF signal (Figure 2F) allows observation of events lasting seconds or minutes, with durations ultimately limited only by photobleaching. A photobleaching event results in a single-step, abrupt change: donor photobleaching reduces both signals to background levels (seen after approximately 90 s in Fig. 2F, magenta arrow), while acceptor photobleaching changes the acceptor signal to background and the donor signal to maximum. Single-step photobleaching confirms that the trace corresponds to a proteoliposome containing a single copy of SecY_MK_EG labelled with one copy of each dye. FRET efficiencies are computed for each time point (up to the photobleaching event) for many traces and collated into a histogram (Figure 2G), which again exhibits a bimodal distribution with E_FRET_~0.2 and ~0.4.

Both methods thus yield almost identical histograms that show co-existence of two states with E_FRET_~0.2 and ~0.4-0.6, which in turn are comparable to the values predicted for the open and closed states (Figure 1S1F), respectively. The similarity of the histograms obtained by the two methods also suggests no bias in the representation of each state under steady-state conditions, irrespective of whether SecY_MK_EG is immobilised to a surface or placed into a freely diffusing proteoliposome in solution. The approximately equal populations of the open and closed states are a result of the steady state conditions in which complexes spend roughly an equal time waiting for the next initiation event (closed) or are translocating (open). As seen in the example TIRF trace in Fig. 2F, the same complex may undergo multiple turnovers. Since our in vitro experiments were done in the absence of signal peptidase, which *in vivo* liberates the secreted polypeptide from the plasma membrane, this observation suggests that SS can spontaneously diffuse into the membrane and its proteolytic removal is not necessary under those conditions.

To further confirm the assignment of the FRET populations to functional states, two control experiments were performed. SecY_MK_EG alone, which was expected to be closed, was examined using confocal microscopy. Surprisingly, a peak was found in the FRET histogram at E_FRET_~0.35, i.e. lower than expected for the closed state observed in the structures (Figure 2S1A). However, the original translocation steady state reactions (shown in Fig 2) were performed in the presence of pOA, ATP and excess SecA. Hence, another control, containing SecY_MK_EG, 1 µM SecA and 2 mM ATP but without pOA, was examined and indeed the FRET efficiency histogram reproduces that of the low FRET closed state (Figure 2S1B). This suggests that the closed state is only attained within the SecY_MK_EG:SecA complex while the SecY_MK_EG alone is either dynamic being in rapid exchange between open and closed states on the experimental timescale, and/or samples partially open states.

The open state control was prepared by first saturating SecY_MK_EG proteoliposomes with pOA and SecA in the presence of ATP (1mM), then rapidly quenching the translocation reaction by addition of excess AMP-PNP (5 mM) – a non-hydrolysable ATP analogue previously shown to keep the translocation complex intact (Deville et al., 2011). This condition resulted in a FRET distribution with a peak below 0.2 (Figure 2S1C), as expected for a trapped open state.

### Plug opening occurs on the ms timescale and is driven by ATP hydrolysis

Both TIRF and confocal microscopy revealed that SecY_MK_EG occupies discrete closed (inactive) and open (translocating) states under steady state conditions. While the transitions between them are seen as instantaneous within the time resolution of the TIRF method (limited by signal to 0.2 s per frame), it is possible to resolve these events with confocal data collection on the millisecond timescale. Initial data acquisition results in bursts lasting from ~1-10 ms (Figure 3S1A), but this time scale can be further extended by the known behaviour of single particles in dilute solutions. As illustrated in Fig. 2&D, highly diluted diffusing particles are likely to revisit the confocal volume within a short time, while entry of another particle in the same time frame is statistically less probable. Thus, closely-spaced photon bursts are usually due to multiple passages of the same vesicle through the confocal volume – a phenomenon called recurrence (Hoffmann et al., 2011). When the diffusion coefficient of the vesicles, the size and shape of the confocal volume and concentration are taken into account, the probability of recurrence of the same proteoliposome can be estimated (Figure 3S1B) and for our preparations remains high (P > 0.9) up to 80 ms. Therefore, the experimental setup provides a large window in which bursts are likely to be generated by the same labelled SecY_MK_EG as it enters and re-enters the confocal volume.

We exploited this phenomenon to follow transitions during translocation initiation with the help of **R**ecurrence **A**nalysis of **S**ingle **P**articles (RASP, (Hoffmann et al., 2011)). To maximise the number of initiation events observed, translocation reactions were started by addition of sub-saturating ATP (0.1 mM) and measured immediately, allowing pre-steady state data collection. Within the resulting bursts, an initial event is arbitrarily selected and its “initial” FRET efficiency computed (designated E1). If another burst is found at a set delay after the initial event (within the high recurrence probability region of a maximum of 80 ms), then this is used to compute a “final” efficiency E2. This is repeated with many bursts for the same delay time and each pair of FRET values is mapped onto a two-dimensional contour plot (Figure 3A). Complexes that remain in the same state appear as spots along the diagonal (as seen in the top panel of Figure 3A), while spots off the diagonal (see Figure 3A middle and bottom) represent state transitions. As the time between E1 and E2 increases, the probability of a state change increases. This is illustrated in Figure 3A (and movie 3S2): after 1.6 ms (top panel), most of the bursts both start and end with the plug closed (E_FRET_~0.4), while as time increases (middle and bottom panels) progressively more complexes transition from closed to open (i.e. appear below the diagonal). The population of states along the E2 axis between the two main spots may be a result of averaging (burst duration being on the same time scale as the transition kinetics) or, alternatively, represent *bona fide* intermediates.

**Figure 3:**
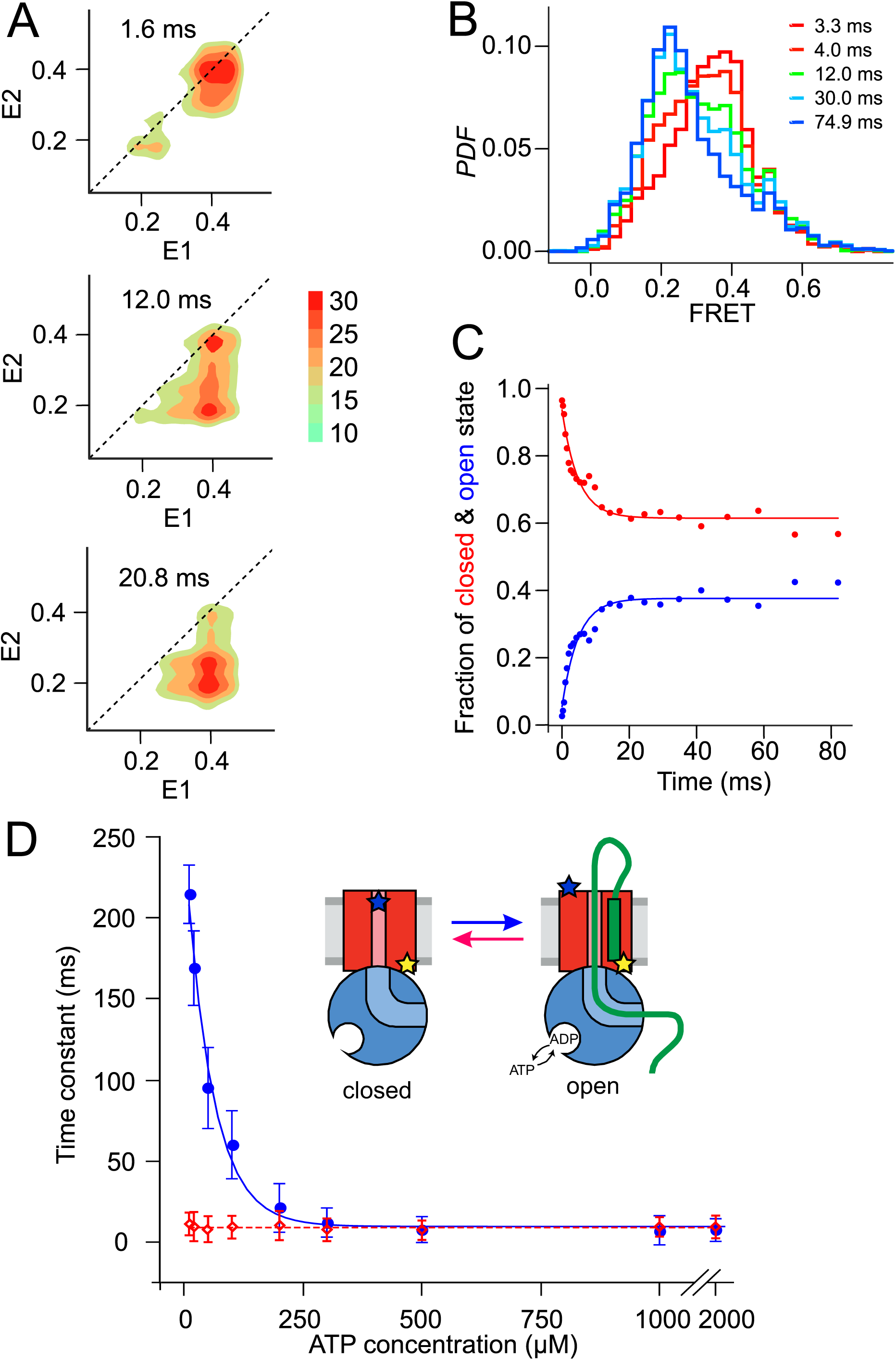
Monitoring plug opening during initiation. **A:** Two dimensional FRET efficiency contour plots (transition density plots) were obtained from bursts collected for SecY_MK_EG:SecA:SecB:pOA in the presence of 0.1 mM ATP using the confocal ALEX setup. The events were classified according to initial FRET (E1) and the burst recurrence FRET (E2) observed after the indicated time delay. Data shown as Probability Density Function (PDF) contour plots with scale on the right. **B:** RASP analysis was performed with the initial state interval centered at 0.4 (± 0.1). RASP PDF shows rapid decreasing closed state population (red) and concomitantly increasing open state (blue). Time in these RASP histograms is colour coded according to the legend. **C:** Opening (blue) and closing (red) kinetic profiles extracted by a two-state approximation. **D:** ATP concentration dependence of the opening (blue) and closing (red) time constants. Open and closed state interconversion is shown schematically in the centre.

The RASP approach effectively allows kinetic rate constants to be determined without the need to synchronise the sample at the single molecule level. As long as the experimental conditions assure that the transitions of interest are taking place, interconversion between different states can be quantified by collating information from many single molecule events. One way to do this is to select initial bursts with a specific E1 value, and follow the time evolution of E2. For example, to monitor plug opening, we selected bursts with starting E_FRET_ values within a window of 0.3-0.5 (closed; centred at the E1~0.4) then plotted E2 after different times in a one-dimensional histogram (Figure 3B). At each time point, the histogram is well described by a two-state model, allowing the open and closed populations to be determined. Fitting these populations as function of time yields the rate of plug opening (Figure 3C). The rate of plug closing can be determined in the same way; both occur on a millisecond time scale.

The dependence of these transition times on the concentration of ATP (or of non-hydrolysable ATP analogues) was next investigated to reveal whether plug opening or closing depend on ATP binding and/or hydrolysis. Figure 3D shows that the opening time constant in the presence of SecA, SecB and pOA increases with decreasing ATP concentration, while closing happens on a ~10 ms timescale and does not depend on ATP. The opening time constant also converges to 10 ms in a saturating ATP concentration. The apparent concentration at which the opening process is at its half maximum rate, *K*_50%_ ~56 µM, obtained by fitting these data, is similar that of ATPase *K*_M_ ~50 µM (Robson et al., 2009a), suggesting that ATP hydrolysis is required for the initial plug opening. Without pOA there are only rare spontaneous opening and closing events, with time constants that do not depend on ATP concentration (Figure 3S3). In the presence of AMP-PNP, SecY_MK_EG remains in the closed state (Figure 3S4). Therefore, ATP binding and hydrolysis together with pre-protein substrate engagement are needed to fully displace the plug from the channel.

We also examined temperature dependence of the opening and closing, over the temperature range 15–37 ºC (288.15- 310.15 K) for which the membrane remains fluid and SecY_MK_EG is active (Figure 3S5). The activation energy for opening, E_a_ ~61 kJ/mol, is close to the value measured for SecA ATPase activity in the presence of translocating substrate or signal peptide (66 kJ/mol), but much lower than that of SecY_MK_EG:SecA alone (~180 kJ/mol) (Gouridis et al., 2009). This suggests that the ATP-driven plug opening is performed by SecA in complex with SecY_MK_EG pre-activated by the SS. On the other hand, the closing transition exhibits significantly lower activation energy – E_a_ ~45 kJ/mol, which is well below values reported for ATPases (60 - 70 kJ/mol) (Jenkins et al., 1999), consistent with closure being independent of ATP (Figure 3D).

### Translocon unlocking by the signal sequence is necessary, but not sufficient, for plug opening

We next used RASP analysis, as described above, to further probe the determinants of plug motion. SecY_MK_EG alone shows considerable static heterogeneity (spread along the diagonal) and dynamics (off-diagonal spots), with a broad diffuse spot between the closed and open configurations (Figure 4A). By contrast, addition of SecA to SecY_MK_EG results in the formation of a much more stable complex, with the plug predominantly residing in the closed state (Figure 4B). Assembling an active translocation complex through the addition of pOA, SecB and saturating ATP to SecY_MK_EG:SecA, then allowing it to reach steady state (>5 min) causes the plug to shift into a predominantly open state, with a hint of some closing transitions represented by a smear along the E1 axis (Figure 4C). This is distinct from the dynamic behaviour of the same complex in the presence of a sub-saturating ATP concentration (0.1 mM; Figure 4D): under these conditions we observe a more dynamic landscape of FRET with significant population of the closed state and transitions to the open state (off-diagonal smear along E2 axis). This further corroborates that plug opening is dependent on ATP concentration.

**Figure 4:**
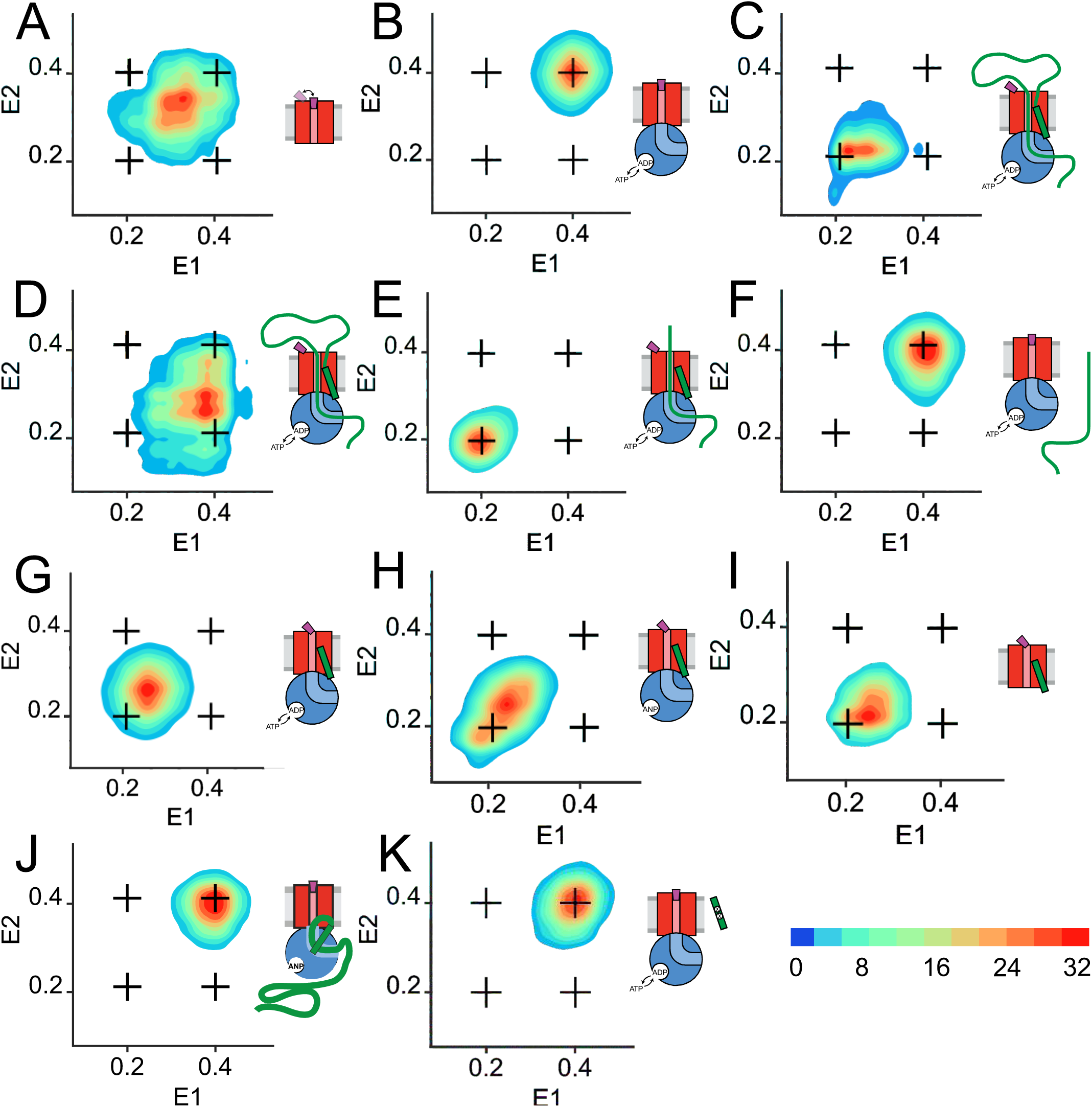
Two-dimensional FRET efficiency histograms (transition density plots). **A:** SecY_MK_EG alone. Transition density for delays up to 21 ms were obtained from RASP analysis of 10,000 events. In all panels the crosshair symbols indicate positions of the open and closed state FRET values within the E1-E2 plot. A scale bar for count contour levels is shown in the lower right corner. A cartoon in each panel schematically depicts the composition of the Sec complex and reaction conditions. **B:** SecY_MK_EG:SecA in the presence of 1 mM ATP. **C:** SecY_MK_EG:SecA:SecB:pOA in the presence of 1 mM ATP. **D:** SecY_MK_EG:SecA:SecB:pOA in the presence of 0.1 mM ATP. **E:** SecY_MK_EG:SecA:SecB:OmpA (no SS) in the presence of SS peptide and 1 mM ATP. **F:** SecY_MK_EG:SecA:SecB:OmpA in the presence of 1mM ATP. **G:** SecY_MK_EG:SecA:SecB in the presence of SS peptide and 1 mM ATP. **H:** SecY_MK_EG:SecA:SecB in the presence of SS peptide and 1mM AMP-PNP (depicted as ANP in the cartoon). **I:** SecY_MK_EG in the presence of SS. **J:** SecY_MK_EG:SecA:SecB:pOA in the presence of 1 mM AMP-PNP (ANP in the cartoon). **K:** SecY_MK_EG:SecA:SecB in the presence of defective (four residue deletion) SS peptide and 1 mM ATP.

It has been shown previously that the SS can unlock the translocon *in trans* and initiate the translocation of mature substrates, *i.e.* those lacking a SS (Gouridis et al., 2009). Structural studies demonstrated that this unlocking process involves SS intercalation within the SecY lateral gate (LG, TMH2 and TMH7, Figure 1) – a process that can occur even in the absence of SecA (Hizlan et al., 2012). Unlocking has also been shown to involve conformational changes at the cytoplasmic side interface with SecA (Corey et al., 2016b; Hizlan et al., 2012), however, the effect of SS binding on the plug location have remained uncertain.

To test the effects of SS addition *in trans* we utilized a synthetic pOA SS (see Methods) and the mature region of pOA (OmpA; i.e. pOA lacking a SS). Addition of SS and OmpA to SecY_MK_EG:SecA in the presence of 1 mM ATP results in a FRET landscape (Figure 4E) akin to that observed for SecY_MK_EG:SecA with pOA (Figure 4C), confirming that SS added *in trans* is indeed capable of stimulating translocation. As a control, we performed the same experiment in the absence of SS and the plug reverted to a closed state (Figure 4F).

Interestingly, when SS is added to SecY_MK_EG:SecA in the absence of OmpA (Figure 4G,H) – even with SecA also absent (Figure 4I) – the plug assumes a new, static state with E_FRET_ ~ 0.25. This state is clearly distinct from the closed (0.4) and open (0.2) states and is well defined with respect to the more dynamic SecY_MK_EG alone (Figure 4A). We assign this newly observed intermediate state as the ‘unlocked’ configuration of the translocon. This observation suggests that binding of the SS peptide is sufficient to unlock the complex (E_FRET_ ~ 0.25) while full opening (E_FRET_ ~ 0.2) is only achieved in the presence of complete protein substrate and ATP hydrolysis by SecA. Indeed, in the presence of non-hydrolysable AMP-PNP SecY_MK_EG:SecA:SecB:pOA remains fully closed (Figure 4J), as is the case when the SecY_MK_EG:SecA:SecB complex is presented with a defective SS (a critical four residue deletion in the hydrophobic section pOA_∆IAIA7-10_ (Emr et al., 1980) (Figure 4K). This confirms that the nature of the SS is critical for the formation of the unlocked complex and suggests that ATP hydrolysis may also be needed to translocate part of the pre-protein and present SS to the LG.

### Translocation rate and slow post-initiation stage

If the low FRET state observed in the TIRF FRET traces (*E_FRET_~ 0.2*, Figure 2F) represents the SecY_MK_EG opening while the substrate is being translocated, then the duration of this state should increase with the length of the pre-protein substrate. Therefore, pOA-based constructs were prepared with various lengths ranging from 100 to 683 aa (where pOA itself is 354 aa; Figure 5). All of these substrates are transported successfully into vesicles as shown by ensemble translocation assays (Figure 5S1) and by the stimulation of the ATPase activity of SecA (Figure 5S2).

**Figure 5:**
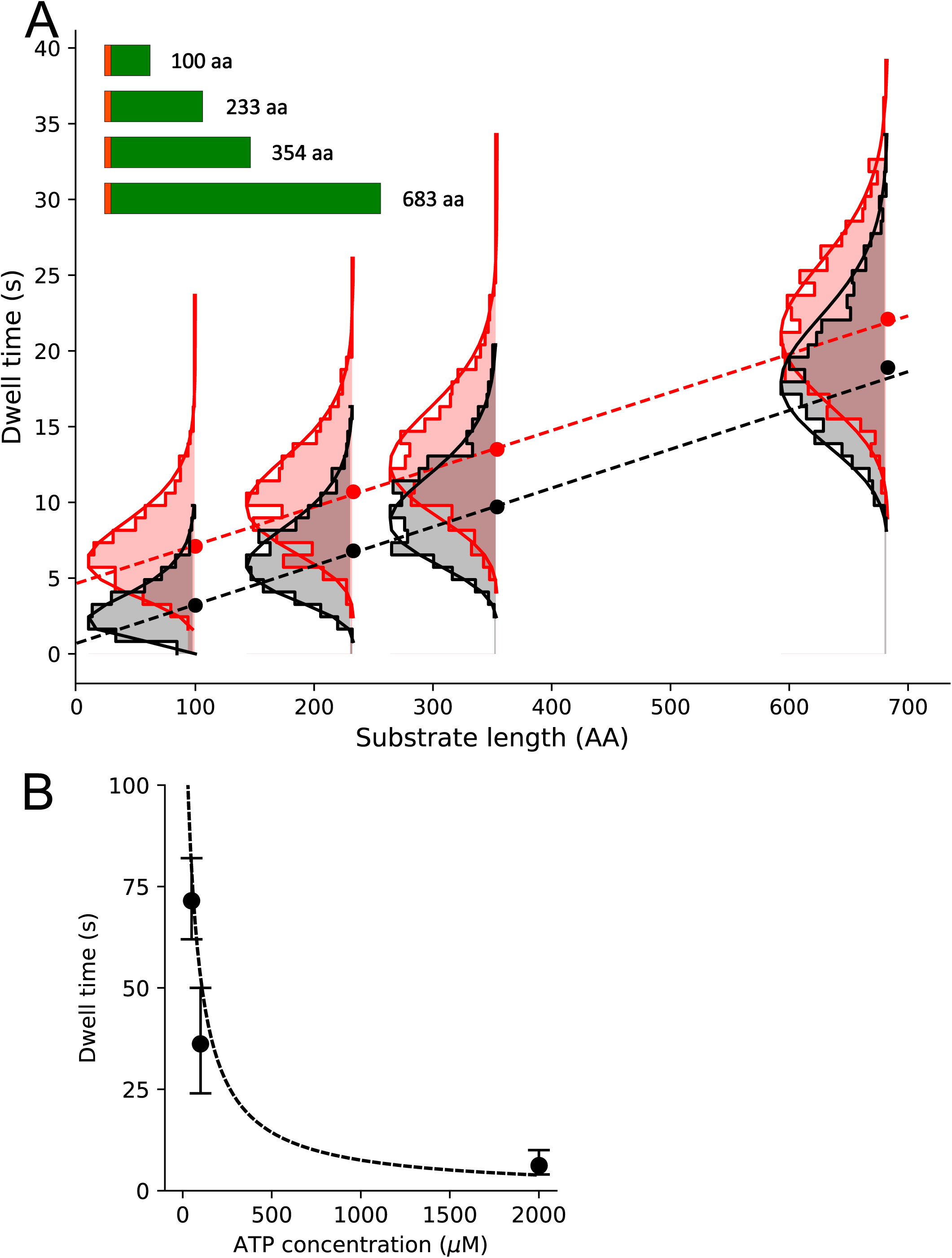
Substrate length dependence of dwell times. **A:** Dwell time dependence for open state (E_FRET_~0.2) on the substrate length (schematically shown in the inset, SS depicted as orange bar) in the presence (black) or absence (red) of SecB. Ordinary least squares (OLS) linear regression (dashed lines) on all photobleaching corrected dwell times (see Figure 5S3 for comparison of uncorrected and corrected dwell time distributions) gave slopes corresponding to translocation rates of 39.6±6.0 aa/s (± standard error) in the absence of SecB and 39.0±6.2 aa/s in the presence of SecB. OLS analysis of the sample with SecB resulted in an intercept close to zero (0.5±0.3 s) while in the absence of SecB the intercept is approximately five seconds (4.7±0.3 s). Overlaid are photobleaching corrected dwell time histograms with gamma function fits (solid lines). Only the distributions for the longest substrate were significantly affected by photobleaching (see Supplementary Figure 5S3 for comparison and Methods for description of the deconvolution correction). **B:** Average open state dwell time dependence on ATP concentration for the shortest 100 aa pOA substrate in the presence of SecB. Error bars were obtained from repeated experiments. The dashed line represents a steady state model with KM fixed at 50 µM and an amplitude scaled to the data (note that paucity of data does not warrant to perform a statistically sound fit).

TIRF traces were obtained for the pOA length variants, and the dwell times of the open state for each were extracted and collated into histograms (Figure 5A). As expected for a stochastic process, a broad distribution of dwell times is obtained for each substrate length. However, the dwell times of the open state clearly increases with the length of the substrate (Figure 5A). By contrast, a similar analysis for the closed state (*E_FRET_ ~0.4*) shows no dependence of the dwell time on the length of polypeptide chain (Figure 5S4). This is not surprising: the closed state dwell time represents an average wait prior to, or between, translocation events and thus reflects the rate of assembly of the active complexes. The formation of the translocation complex would be expected to depend on the concentrations of individual components and other reaction conditions, which were kept constant throughout the experiments.

If the dwell time of the open state is indeed measuring translocation time, then it should increase at sub-saturating ATP concentrations. As shown in Figure 5B, this is indeed the case, and the apparent *K*_M_ (~50 µM) matches that of SecA ATPase (Robson et al., 2009b). Together with the pre-protein length dependence and the short opening and closing times (see above), this justifies the use of the open state dwell times as a proxy for the duration of individual translocation events. These durations can then be combined to estimate the rate at which the polypeptide is being translocated.

To extract the translocation rate per residue, the dwell times were plotted as a function of the substrate length and the slope was obtained using a linear fit (ordinary least squares method using all dwell times, Figure 5A). Assuming a linear dependence of the open state dwell time on the substrate length, the slope yields the average time necessary to translocate a single amino acid. The resulting rate is ~ 40±6 aa/s and within statistical error is independent of cytoplasmic chaperone SecB (Figure 5A).

An extrapolation of the least squares fit line to zero substrate length suggests that there is a significant fixed, length-independent dwell time component of ~5 s in the absence of SecB (Figure 5A). This may be related to a slow initial translocation phase after the channel is already opened; alternatively, it could represent the time it takes for the plug to return to the closed state after the translocation event is complete. However, this constant dwell time decreases close to zero in the presence of SecB. As SecB acts on the cytoplasmic side, this suggests this slow phase is related to the delivery of the substrate to SecA – with SecB presumably making the initial handover and delivery of substrate more efficient. This is consistent with the observation that SecB reduces ATP consumption, as indicated by ensemble ATPase assays with SecA (Fig.5S2). It is also unlikely that the translocation channel would remain open for several seconds after translocation: *in vivo* this would dissipate the PMF for no obvious gain. Hence, we assign the constant dwell time to a slow initial translocation phase, which follows ATP-driven opening of the plug and is accelerated by SecB.

## Discussion

While structural analysis provides exquisite details of characterized intermediates in a protein reaction cycle it is often difficult to position these snapshots in the right order along the reaction coordinate, especially when dealing with multi-subunit assemblies and complex substrates like biopolymers. Likewise, it is hard to detect and structurally characterize intermediates in complex reactions that cannot be synchronized. Single molecule techniques are able to overcome both of these problems, providing high sensitivity, time-resolved detection of otherwise elusive intermediates, without the need for a synchronized reaction.

Here, we have combined two complementary single molecule fluorescence techniques to examine the process of pre-protein translocation through the SecY_MK_EG translocon on timescales ranging from milliseconds to minutes. A judiciously placed pair of reporter dyes on the SecY plug and a cytoplasmic reference site allowed us to exploit FRET changes to monitor opening and closing of the channel and to detect intermediates in the translocation process. We can now present a full circle of events and intermediates starting from the resting state *via* the already known unlocked translocon (Hizlan et al., 2012) through a new, ATP-dependent plug opening, followed by slow and fast translocation phases and closure (Figure 6).

**Figure 6:**
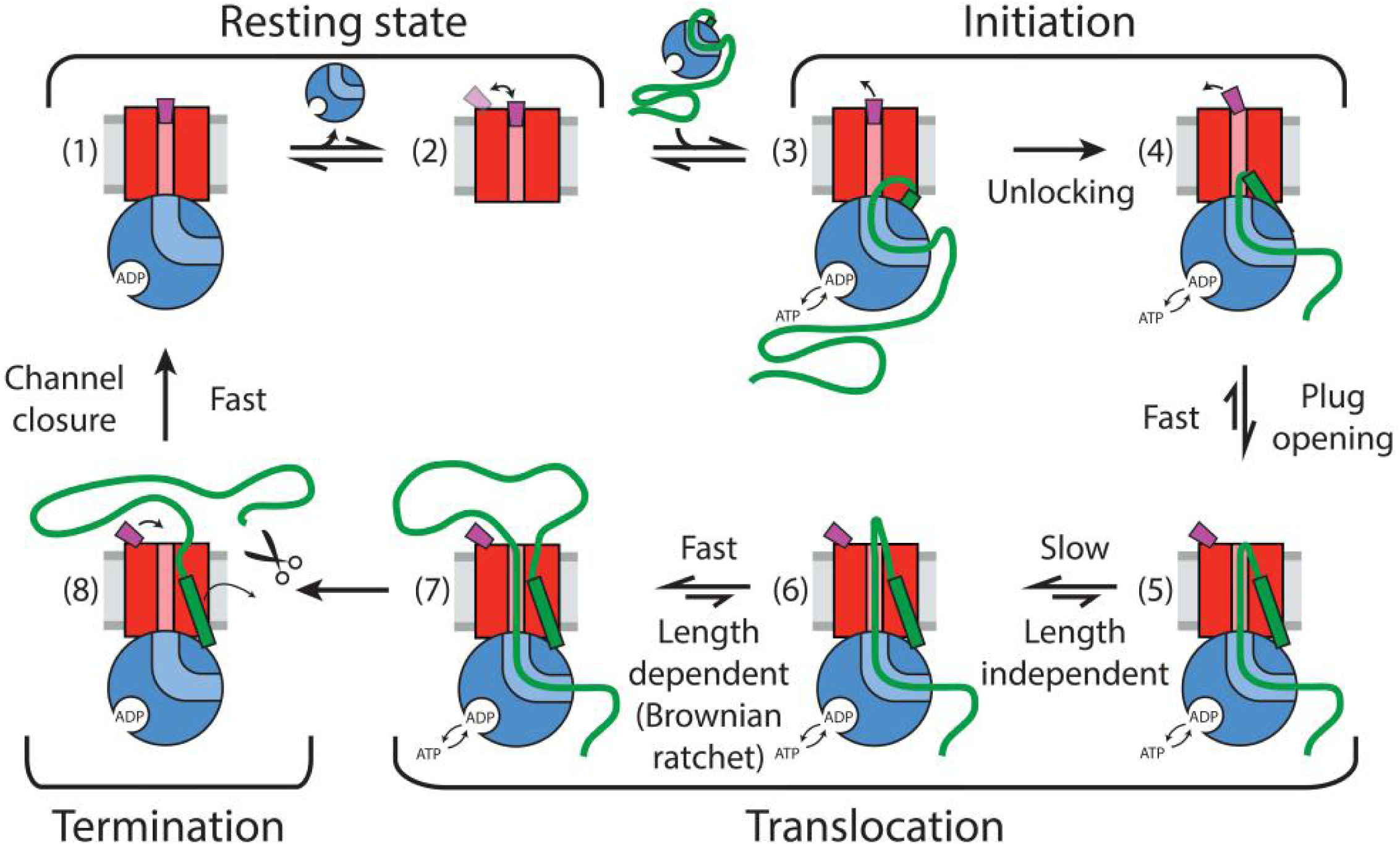
Summary of detected plug states and initiation stages. Colour coding: SecYEG-red, SecA – blue, pOA/OmpA – thick green line, SS –green rectangle, stationary plug – purple, plug in intermediate or transient state – white, lipid bilayer – grey. Scissor symbol indicates substrate liberation by signal peptidase (if present). Thin arrows indicate motion of plug and SS.

The single molecule analysis revealed considerable dynamics of the plug within the resting translocon (state (2) in Figure 6); perhaps consistent with observed variations of its position within SecY_MK_EG. RASP analysis revealed spontaneous fluctuations on the millisecond time scale as evidenced by a broad range of states accessible (Fig. 4A), but showed a more stable closed state when bound to SecA (Fig. 4B). This might help to keep the channel sealed when activated and partially opened by SecA, but in the absence of a substrate (state (1) in Figure 6).

Biochemical and structural methods have described an activation, or ‘unlocking’, of the protein channel by the SS (Corey et al.; Gouridis et al., 2009; Hizlan et al.). We show here that the SS alone is sufficient to partially displace the plug, but not to fully open the channel. For this, the SS and the succeeding stretches of the mature polypeptide chain need to engage the channel, in conjunction with SecA. Crucially, we show that this state also requires ATP hydrolysis when SS is presented as part of pre-protein (states (3) and (4) in Figure 6).

The ATP dependence of the rate of channel opening (KM ~50 µM) and its apparent activation energy (61 kJ/mol) mirror that of the activated SecA ATPase (Gouridis et al., 2009; Robson et al., 2009b). These combined observations suggest that the rate limiting steps of channel opening and the hydrolytic cycle of ATP are identical. The simplest explanation is that the initial rate limiting step is the ATP driven intercalation of pre-protein into the channel, forcing the SS and N-terminal hairpin of the mature sequence towards the periplasm with concomitant displacement of the plug (transition (4) to (5) in Figure 6).

Following the fast channel opening and substrate intercalation, a slow (up to 5 s in the absence of SecB) initiation phase, independent of substrate length takes place (transition (5) to (6) in Figure 6). During this phase, the plug is already opened but translocation might be arrested or is very slow. This phase could be associated with a slow conformational re-arrangement of the translocon, *e.g.* repositioning of the SS within the LG or threading of the initial loop more completely through the translocon. Given that the duration of this phase is reduced significantly by SecB on the cytoplasmic side of the membrane, we speculate that the slow phase might be related to the unfolding of residual structure in the polypeptide substrate by the SecA:SecB complex, making the initial delivery of the substrate to SecA more efficient. In the absence of SecB this might require multiple rounds of ATP hydrolysis with little translocation achieved. This is further supported by the higher consumption of ATP by short polypeptide substrates, the increased efficiency for such substrates in the presence of SecB (Figure 5S2) and the previously observed compensatory genetic link between the SecA and SecB genes (Cook and Kumamoto, 1999).

The slow phase is followed by a substrate chain length-dependent phase, which occurs at an average rate of ~40 aa/s (state (6) to (7) in Figure 6). For each length of substrate, the distributions of rates vary considerably (Figure 5A), as expected for a stochastic process which involves long heterogeneous substrates, consistent with previously proposed stochastic translocation mechanisms (Allen et al., 2016; Bauer et al., 2014). The rate of processive translocation is higher than the previously reported value of ~ 30 aa/s which were also estimated using pOA substrates with various chain lengths, but in the absence of SecB (De Keyzer et al., 2002; Tomkiewicz et al., 2006). The difference is readily explained by the slow initiation phase which is substantial only in the absence of SecB. In the presence of SecB the rate estimated here would support transport of typical individual precursors (100 to 1000 amino acids) within 25 s which is close to the previously estimated transport requirements (~20 s per substrate) in rapidly dividing *E. coli* cells (Brundage et al., 1990).

Channel closure monitored by the relocation of the plug is fast (< 10 ms) and ATP independent. Hence, it does not seem to impose any limitation on the overall translocation efficiency (state (7) to (8) in Figure 6). Evidently, a single translocon is capable of multiple rounds of translocation, at least for pOA, even in the absence of signal peptidase. This suggests that the signal peptide together with the rest of the substrate is able to diffuse from the translocon and does not perturb plug closure (state (8) to (1) in Figure 6).

In summary, we have devised a novel single molecule assay that allowed the determination of the intrinsic rate of polypeptide translocation for the first time. We have also characterised previously known steps in the initiation of translocation and discovered new stages in the reaction, all of which were mapped onto the overall cycle of translocation. Together the results provide a refined framework for understanding the molecular mechanism of ATP driven protein translocation that draws on the powers of single molecule measurements to unpick such complex reaction mechanism in unsynchronised systems.

## Methods

### Protein preparation

Site-directed mutagenesis was performed using the QuikChange protocol (Agilent) and confirmed by sequencing. SecY_MK_EG, SecA, SecB and full length pOA were produced as described previously (Deville et al., 2011; Gold et al., 2007; Whitehouse et al., 2012). Different lengths of pOA were produced adopting existing methods (De Keyzer et al., 2002). OmpA lacking the SS was purified as described in (Schiffrin et al., 2016)). SecY_MK_EG was produced in the same way as wild-type, then labelled for 45 mins on ice at 50 µM with 100 µM each of Alexa 488-C5-maleimide and Alexa 594-C5-maleimide (Invitrogen). The reactions were quenched with 10 mM DTT, and excess dye removed by gel filtration (Superdex-200, GE Healthcare, UK). Labelling efficiencies were between 75–90% for each dye, as determined using the manufacturer’s quantification method and assuming a molar extinction coefficient of 70,820 cm^−1^ for SecY_MK_EG.

### SmFRET in msALEX TIRF configuration on immobilized proteoliposomes

SecY_MK_EG was reconstituted into proteoliposomes (PLs) with *E. coli* polar lipid to a final concentration of 1.5 nM, and extruded to 100 nm: at this concentration and size, most liposomes are expected to contain either 0 or 1 copy of SecY_MK_EG (Deville et al., 2011).

PLs were immobilized on a glass supported lipid bilayer and imaged with a previously described TIRF set-up (Sharma et al., 2014) extended with msALEX illumination (Kapanidis et al., 2005). The alternation cycle consisted of 100 ms cyan (488 nm) and orange (594 nm) excitation periods, adding information about stoichiometry of dyes and thus allowed to filter out singly labelled molecules.

The buffer used was TKM (20 mM Tris pH 7.5, 50 mM KCl, 2 mM MgCl2) with 1 mM 6-hydroxy-2,5,7,8-tetramethylchroman-2-carboxylic acid (TROLOX) and GODCAT, enzymatic oxygen scavenging system, consisting of a mix of glucose oxidase, catalase, β-D-glucose (Aitken et al., 2008) to limit photobleaching. Immobilized samples were supplemented with 1 µM SecA, 10 µM SecB (if present), 700 nM pOA (if present), 1 mM ADP or 1 mM AMPPNP or varying concentrations of ATP (unless stated otherwise in figure legend). Translocation assays under saturating ATP conditions were supplemented with an ATP regeneration system. TIRF movies (200 ms resolution) were taken from samples immediately after mixing of all components directly at the microscope stage.

The data were analyzed in iSMS software (Preus et al., 2015). The two channels of each image were aligned and fluorescence count traces (donor and acceptor) were extracted and raw FRET efficiencies (E) and stoichiometries (S) were computed. To eliminate contributions from complexes with single type of dye or photobleached acceptor dye, only traces with S values between 0.25 and 0.75 were selected for further analysis. Another selection criterion for molecules was anti-correlation of intensity in donor and acceptor channels. All trajectories were also checked for bleaching and blinking events. Molecules showing bleaching were used to obtain correction factors, i.e. donor leakage, direct acceptor excitation and gamma factor (Preus et al., 2015). Experiments were repeated at least three times using independent proteoliposome preparations. Corrected FRET values were used to produce histograms.

### Photobleaching correction

Statistical distributions of dwell times were corrected for photobleaching using probability distribution of photobleaching times (P_photobleaching_) estimated from TIRF traces for each experiment. Subsequently we employed non-negative regularized iterative reconvolution of two distributions, i.e. photobleaching of individual molecules and simulated dwell time distribution to match measured data:

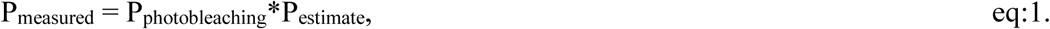

where P_measured_ relates to the Probability Density Function (PDF) derived from experiment, P_photobleaching_ is PDF of photobleaching (estimated from data) and P_estimate_ is the simulated gamma distribution with scale equal to one, representing an estimate of the original data unaffected by photobleaching. The best hit was found by a least squares method as implemented in SciPy optimize python package (http://www.scipy.org/). A regularized inverse transform was used to reconstruct dwell-time histograms using non-negativity constraints (Provencher, 1982) and the reconstructed probability density function, P_estimate_.

### µsALEX confocal experiments on freely diffusing proteoliposomes

The experimental set-up used to collect µsALEX data was previously described (Sharma et al., 2014). The laser alternation period was set to 40 μs (duty cycle of 40%) with intensity for the 488-nm laser ∼100 µW and the 594-nm laser intensity ∼90 μW. Data were collected using Labview graphical environment (LabView™ 7.1 Professional Development System for Windows, National Instruments, Austin TX, USA) (Lee et al., 2005). Separate photon streams were then converted and stored in an open file format for timestamp-based single-molecule fluorescence experiments (photon-hdf5), which is compatible with many recent data processing environments (Ingargiola et al., 2016a).

Fluorescence bursts were analyzed by customized python programming scripts based on the open source toolkit for analysis of freely-diffusing single-molecule FRET bursts (Ingargiola et al., 2016b). The background was estimated as a function of time, respecting exponentially distributed photon delays generated by a Poisson process. In order to guarantee a maximal signal-to-background ratio, we used background dependent dual-channel burst search (DCBS) (Michalet et al., 2013; Nir et al., 2006) in sliding window mode (Eggeling et al., 1998), which effectively deals with artifacts due to photophysical effects such as blinking. Further filtering was based on dye stoichiometry (S within 0.25-0.75).

Three correction parameters: γ-factor, donor leakage into the acceptor channel and acceptor direct excitation by the donor excitation laser were employed and determined using polyproline standards of different length as FRET samples (Best et al., 2015; Sharma et al., 2014). Corrections were applied at the population-level (Lee et al., 2005) to avoid distortion of the FRET distributions (Gopich and Szabo, 2007).

Filtered bursts were then assembled into 2D E-S histograms and 1D probability density function plots were generated using library Seaborn, based on Matplotlib (Hunter, 2007). Subpopulations were fitted to weighted Gaussian mixture models using Scikit (Pedregosa et al., 2011).

### Recurrence Analysis of Single Particles (RASP)

To analyze events on timescales from 100 μs to ~ 100 ms, we employed RASP which relies on extremely diluted samples, where the probability for a molecule to return to the confocal volume is greater than the probability of a new molecule being detected (Hoffmann et al., 2011). RASP extracts time resolved information for FRET subpopulations by constructing recurrence FRET efficiency histograms. These are acquired by first selecting photon bursts from a small transfer efficiency range (initial bursts) and then building the FRET efficiency histogram only from bursts detected within a precisely defined short time interval (the recurrence interval) after all selected initial bursts. Systematic variation of the recurrence interval allows determination of the kinetics of interconversion between subpopulations.

The longest usable recurrence time is related by concentration and diffusion time of the observed objects and can be set based on the recurrence probability. To estimate the recurrence probability of single molecules, we employed a correlative approach (Hoffmann et al., 2011). Bursts from different and non-interacting molecules are expected to be uncorrelated. On the other hand bursts originating from the same molecule should be correlated and a “same molecule” probability P_same_(τ) was calculated as:

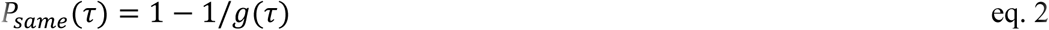

where g(τ) is the burst time autocorrelation function of all detected bursts. From a fit to the data, we determined for each burst pair the probability that it originated from the same, recurring molecule, and calculated the average P_same_ for a subset of bursts by averaging over all corresponding burst pairs.

### Recurrence Transfer Efficiency Histograms

To derive kinetics from RASP, we constructed transfer efficiency histograms from a set of bursts selected by two criteria. First, the bursts b_2_ must be detected during a time interval between t_1_ and t_2_ (the “recurrence interval”, T = (t_1_, t_2_)) after a previous burst b_1_ (the ‘initial burst’). Second, the initial bursts must yield a transfer efficiency, E(b_1_), within a defined range, ΔE_1_ (the ‘initial E range’). The set R of burst pairs {b_1_, b_2_} selected by these criteria is then:

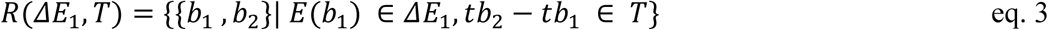

where tb_1_ and tb_2_ are the detection times of the bursts b_1_ and tb_2_, respectively. The set of burst pairs R is the starting point for the different types of analysis presented here. A very informative way of representing the data is the FRET efficiency histogram of all values E(b_2_), the “recurrence transfer efficiency histogram”.

### Cross-peaks in 2D recurrence transfer efficiency contour plots

As a visual guide we constructed 2D transition density contour plots. They were obtained from two-dimensional Gaussian KDE analysis (Scott, 1992) of burst pairs, where the initial burst and the second burst yield transfer efficiencies in range ΔE1 and ΔE2, respectively. Each plot was constructed for a certain recurrence interval T.

2D contour plots were also used to address a common issue in the analysis of transfer efficiency histograms: the determination of the number of contributing subpopulations and their peak shapes. These were answered by choosing short recurrence intervals and initial transfer efficiency ranges that represent only a single subpopulation. The significance of small populations and the properties of strongly overlapping peaks were tested with this approach.

### Interconversion dynamics from kinetic recurrence analysis

To extract rates of interconversion between subpopulations from time-dependent recurrence E histograms, we constructed histograms for different recurrence intervals, and extracted the fraction of a subpopulation versus time. To determine the rates of interconversion, we related the change in the recurrence E histograms with increasing recurrence time to the dynamics of the interconversion process as was first shown by (Hoffmann et al., 2011).

For a system populating two states A and B, we defined the probability pA (τ,ΔE_1_) that from the set of burst pairs R(ΔE_1_,T), b2 originates from a molecule in state A. pA (τ, ΔE_1_) was determined from global fitting the corresponding recurrence histogram and determining the ratio of the peak area corresponding to subpopulation A over the total area under the peaks corresponding to A and B (for details see (Hoffmann et al., 2011)).

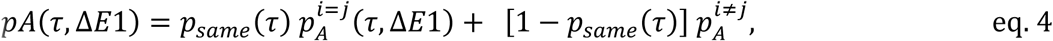

where 
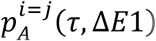
 denotes the probability that a recurring molecule ( *i* = *j*) is in state *A*, and 
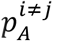
 is the probability that a newly arriving molecule ( *i* ≠ *j*) (leading to burst *b*_2_) is in state 
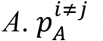
 is probability of measuring a burst originating from a molecule in state A and it was determined from globally fitted areas under the corresponding peak functions extracted from a set of transfer efficiency histograms.

The time dependence of 
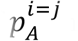
 is determined by the interconversion kinetics between states A and B and is defined as:

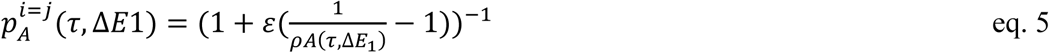

with

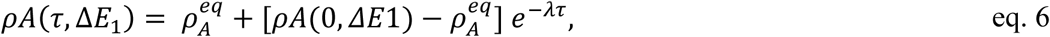

Here, *ρA*(*τ*, ∆*E*_1_) is the probability that a protein that emitted a burst at time 0 with a transfer efficiency in the range Δ*E*_1_ is in state *A* at time *τ*. *ρA*(0,∆*E*_1_) and 
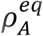
 are the corresponding initial and equilibrium probabilities, respectively. The kinetic rate constant λ corresponds to the sum of the forward and backward rate constants of interconversion between A and B.

## Acknowledgements

This paper is dedicated to our friend and colleague, the late Prof. Steve Baldwin. This work was funded by the BBSRC: TF, RT and SER (BB/N017307/1), DW and IC (BBSRC: BB/N015126/1 and BB/I008675/1), PO, RT, SER and SAB (BB/I008675/1), JEH (BB/M011151/1), IC and WJA (BB/I006737/1); RAC (BBSRC South West Bioscience Doctoral Training Partnership and BB/M003604/I). Additional support was provided by the Wellcome Trust (104632) to IC and WJA and ERC ((FP7/2007-2013) / ERC grant agreement 32240) to SER. TF and RT are supported from European Regional Development Fund-Project “Mechanisms and dynamics of macromolecular complexes: from single molecules to cells” (No. CZ.02.1.01/0.0/0.0/15_003/0000441).

## Supplemental Figure Legends

**Figure 1S1:**
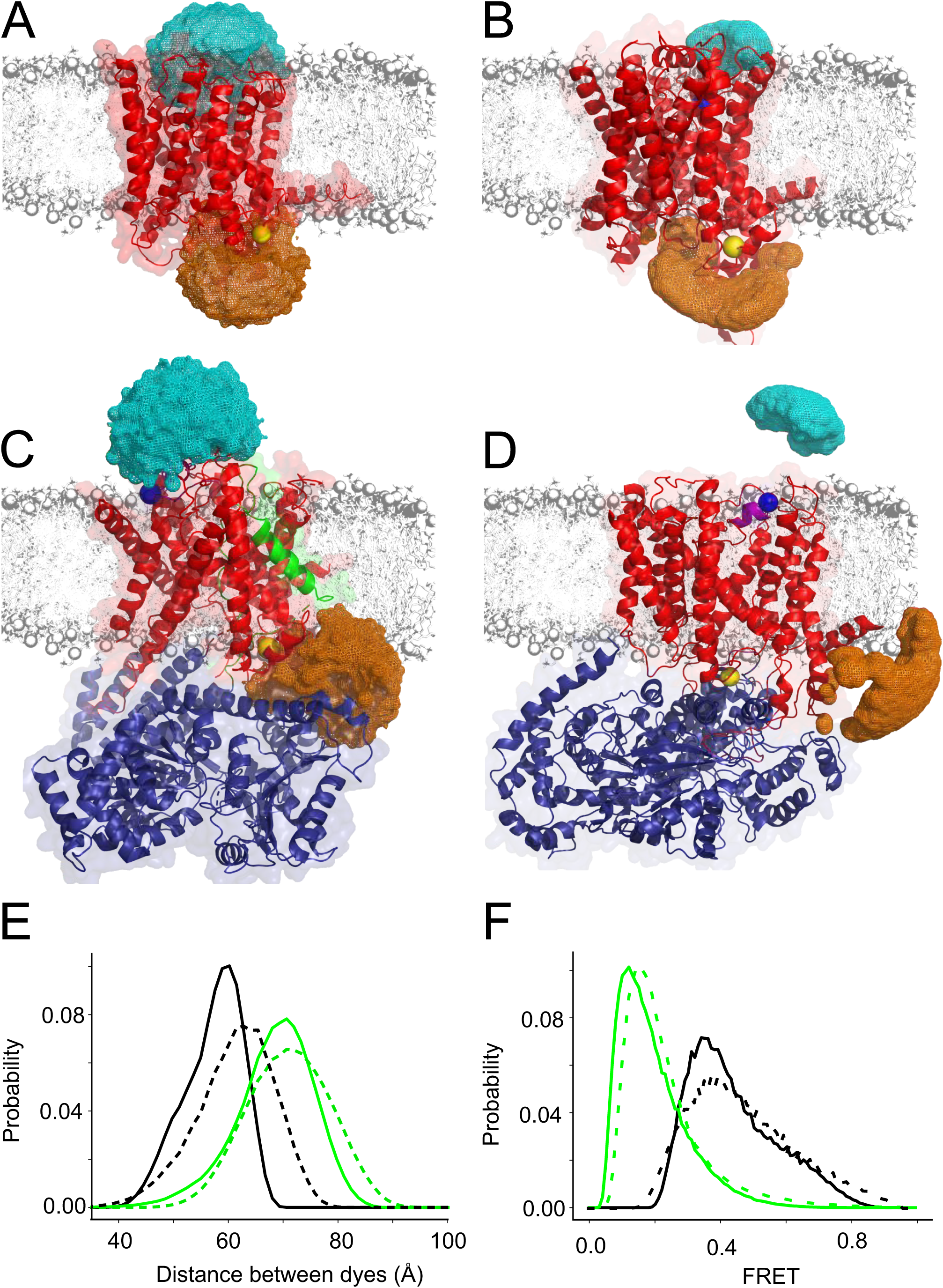
Modelling of fluorescent dye accessible volumes. **A:** Closed state (PDB: 1rhz) (Van den Berg et al., 2004). Accessible volumes (orange for dye attached to the reference residue and cyan for the plug probe) obtained by modelling the dye positions when attached to the mutated residues (blue and yellow spheres) via a short C6 aliphatic linker. Ribbon colours: SecYEG (red), SecA (blue), SS (green). **B:** As **A**, but for the Closed state (PDB:5aww) (Tanaka et al., 2015). **C:** As **A**, but for the Open state (PDB:5eul) (Li et al., 2016). **D:** As **A**, but for the open state (PDB:3din) (Zimmer et al., 2008). **E:** Distributions of inter-probe distances computed from the accessible volumes in panels AD. Open states are in green (PDB: 5eul solid and 3din dashed) while closed states are black (PDB: 5aww solid and 1rhz dashed). **F:** Distribution of FRET efficiencies derived from available volumes. Colour coding as in panel E.

**Figure 1S2:**
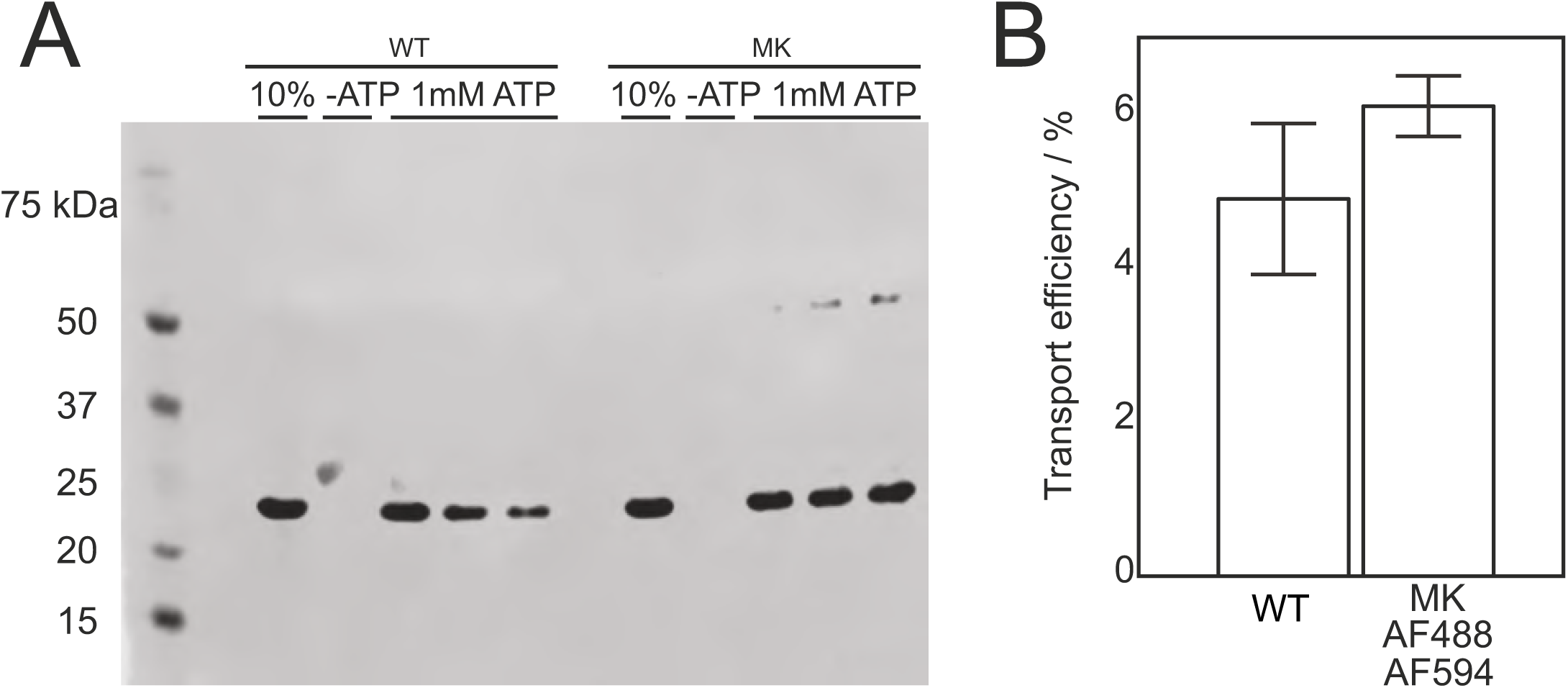
Activity of dual labelled SecY_MK_EG in a translocation assay. **A:** Protease protection assay performed with wild-type (wt) SecYEG and SecY_MK_EG mutant (MK lanes) labelled with AlexaFluor dyes. A 233 amino acid N-terminal fragment of proOmpA was used as substrate. 10% lane corresponds to the 10% fraction of the substrate used and was used to quantify the amount of protected polypeptide after quenching the reaction (lanes labelled 1 mM ATP represent triplicate samples). A negative control reaction was performed in the absence of ATP (-ATP lane). **B:** Efficiency obtained by densitometry using the density of 10% control band as an internal standard.

**Figure 2S1:**
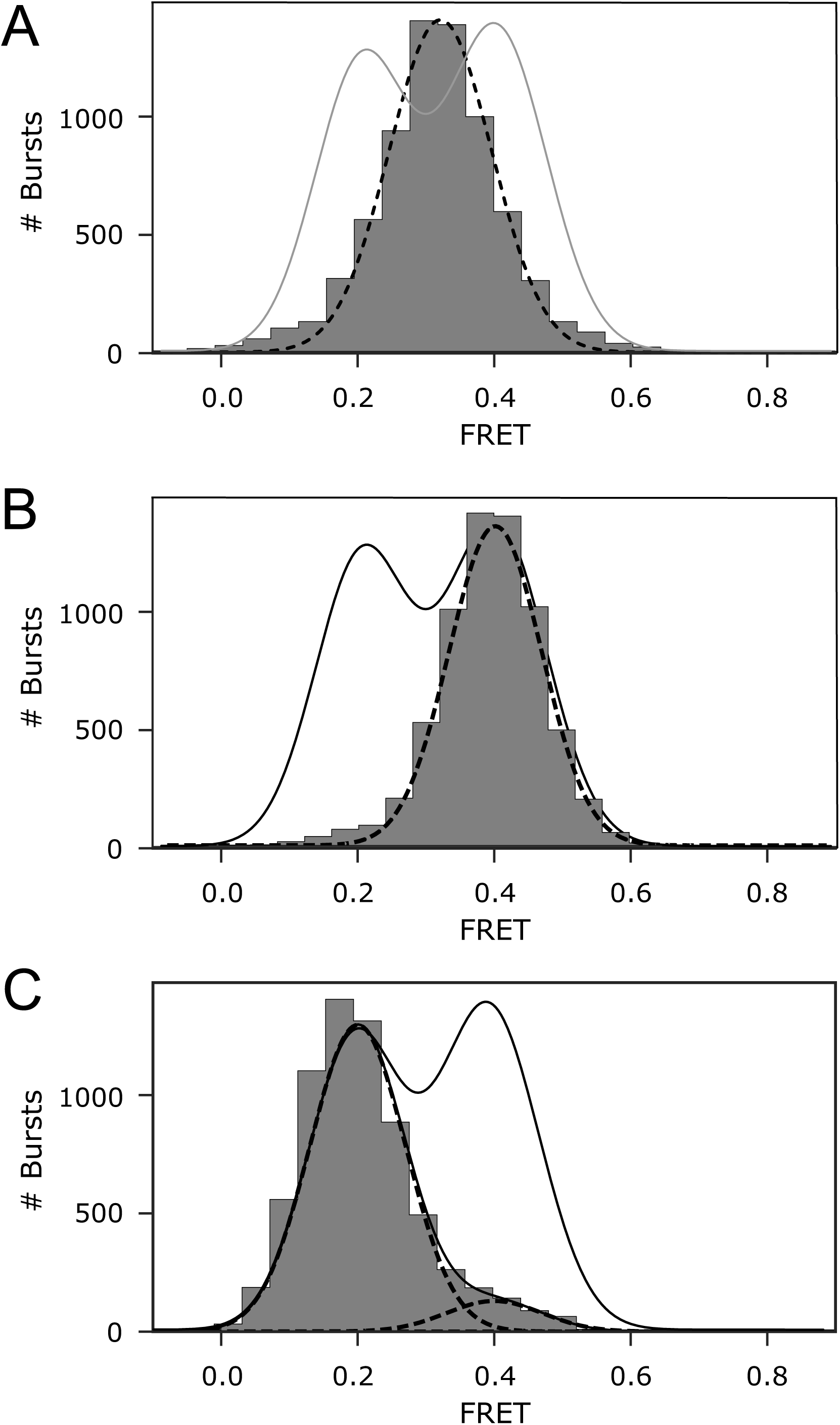
1D FRET efficiency histograms for controls. **A:** SecY_MK_EG alone. All data were collected using the confocal setup and histograms are fitted to a single or sum of multiple Gaussians (dashed) and overlaid with the resulting fit from Figure 2, panel E (grey). **B:** SecY_MK_EG:SecA in 2 mM ATP. The histogram was corrected for contribution from the 50% SecY_MK_EG in opposite orientation which is unable to bind SecA and translocate (unresponsive population), i.e. by subtracting appropriately scaled histogram shown in Panel A from the data. **C:** Open state trapped by addition of 5 mM AMP-PNP to translocating SecY_MK_EG:SecA:SecB:pOA:ATP. Corrected for the contribution from the unresponsive population of SecY_MK_EG as in Panel B.

**Figure 3S1:**
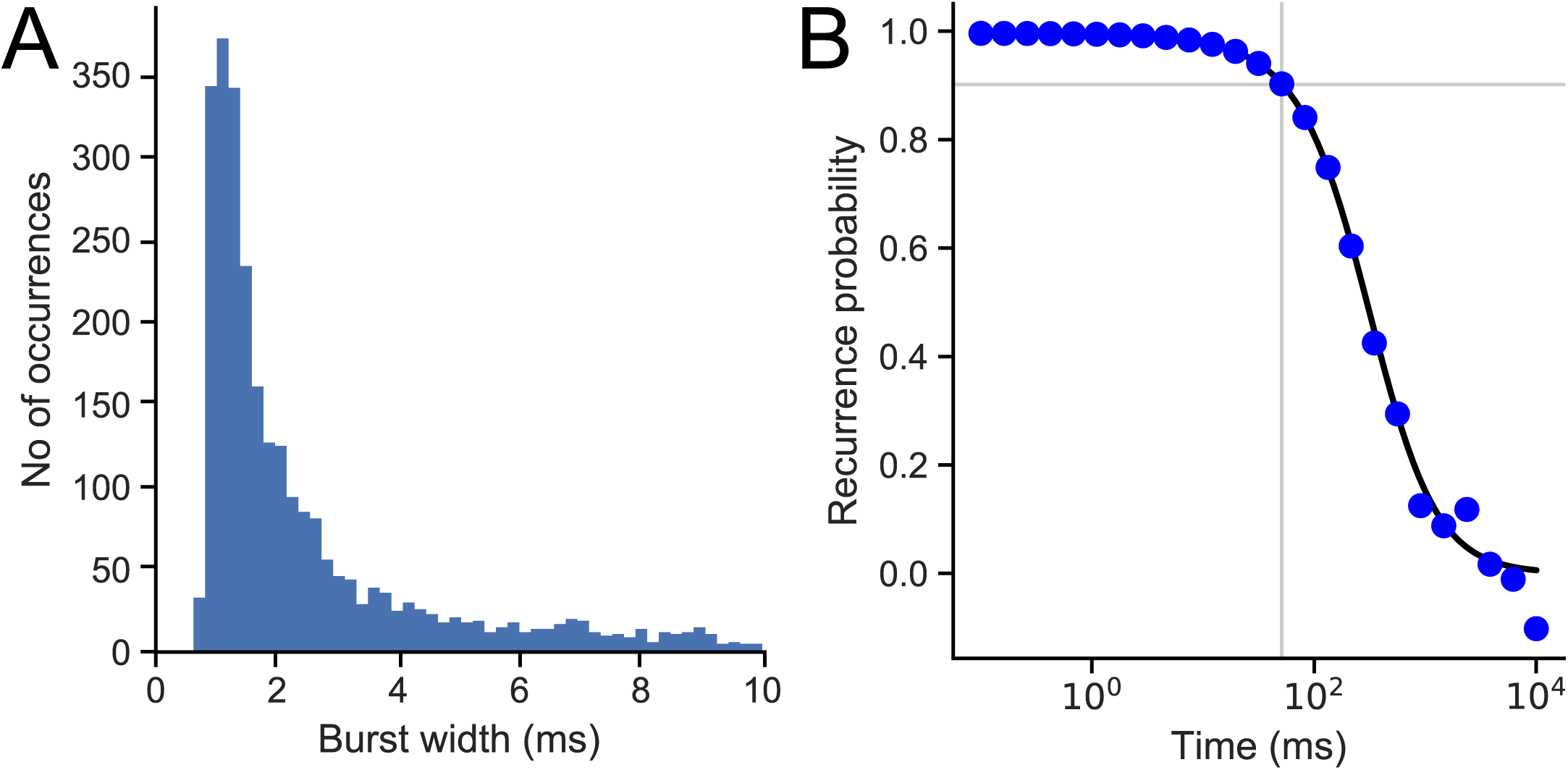
Burst duration and recurrence probability. **A:** Burst width distribution for diffusing proteoliposomes. **B:** Recurrence probability was obtained from data in **A** as described in Methods. The recurrence time of 80 ms, corresponding to the 0.9 probability level, is indicated by grey vertical line.

**Figure 3S2:**
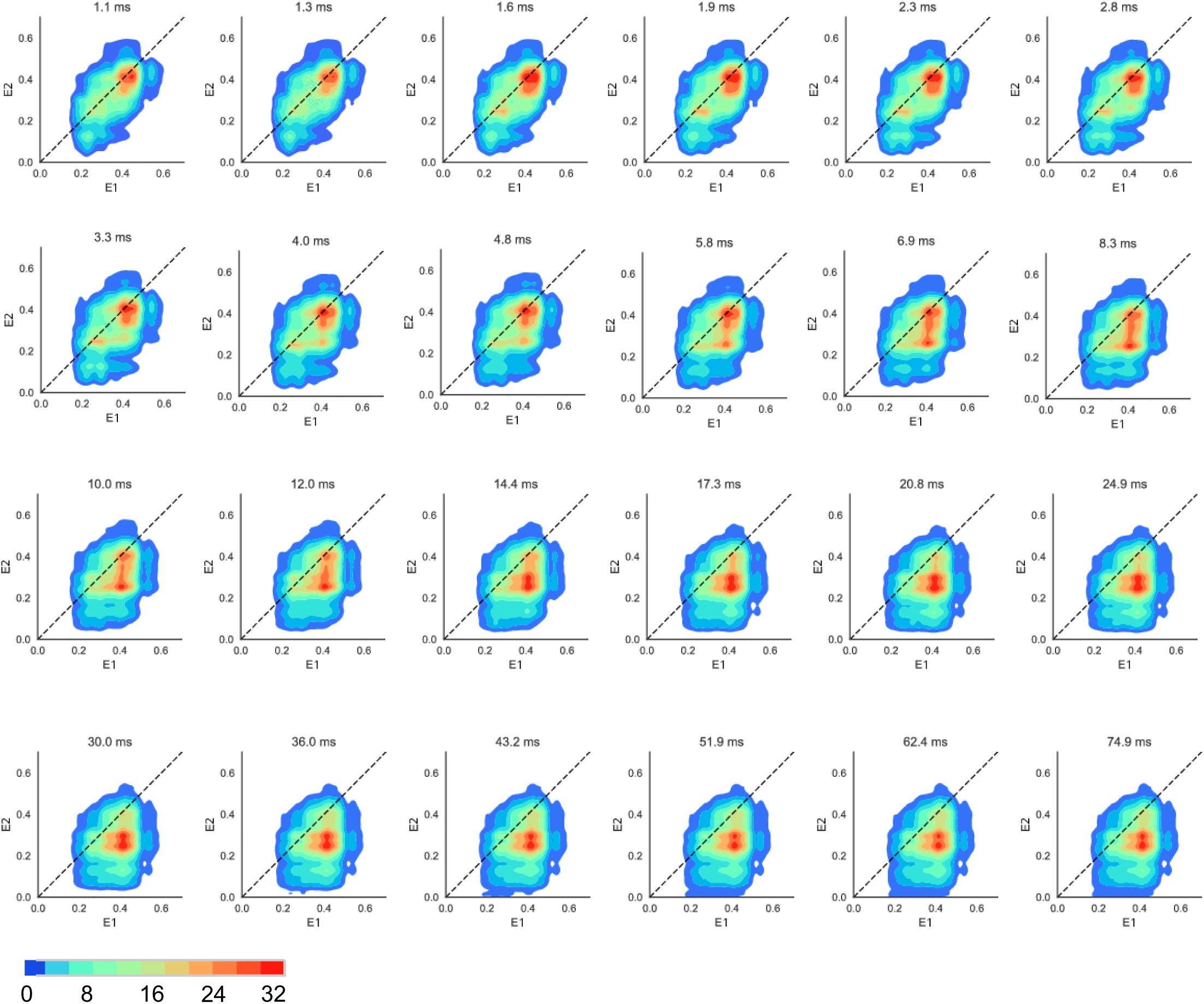
2D transition density plots. A detailed sampling of time resolved transition density plots shown in Figure 3A. Note a wider scale (count contour level bar below) was used compared with Figure 3A. An animation is shown in Supplementary_movie_1 online.

**Figure 3S3:**
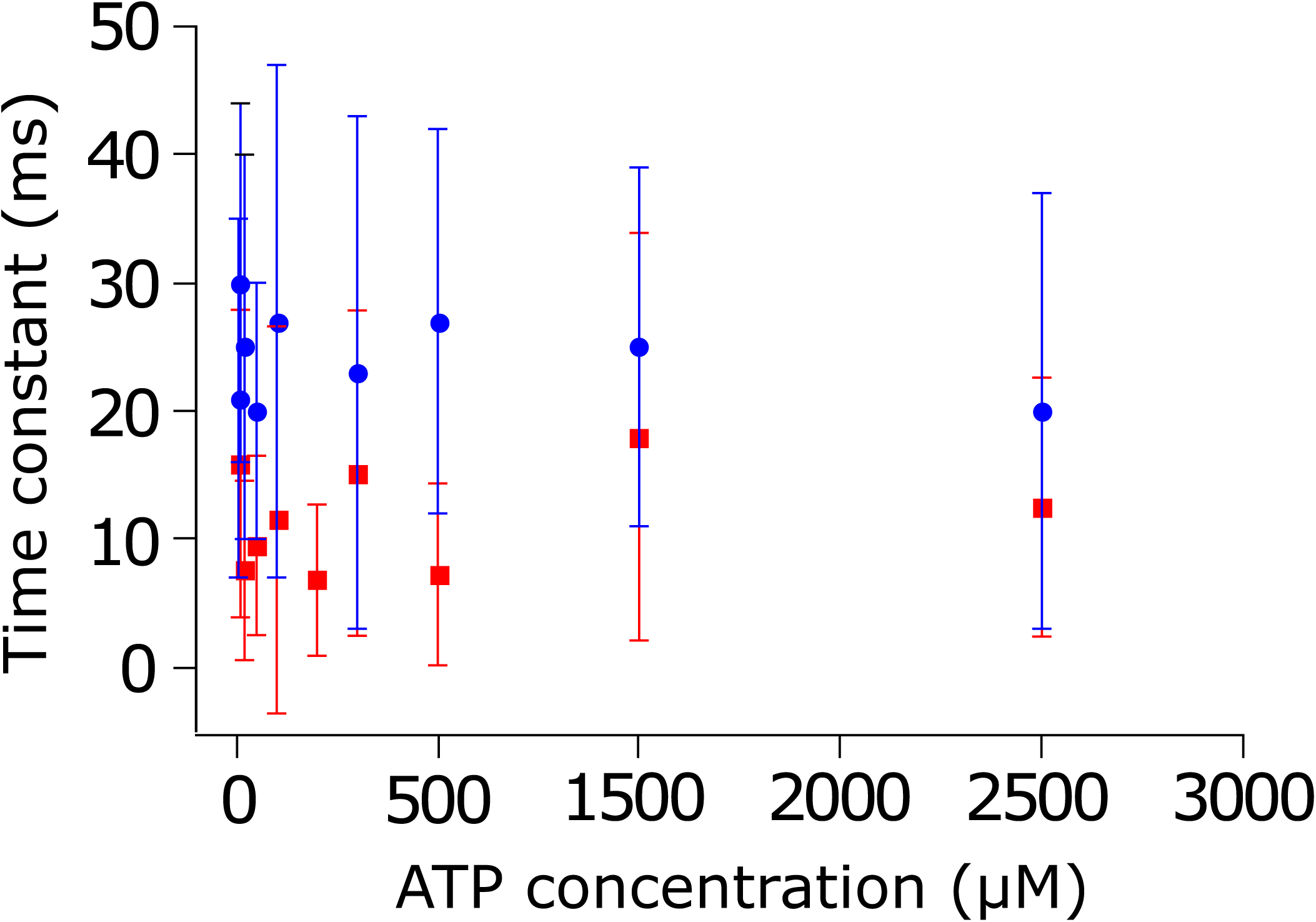
Spontaneous opening and closing in the absence of translocation substrate. Opening (blue) and closing (red) transition times as a function of ATP concentration for SecY_MK_EG:SecA:SecB in the presence of 1 mM ATP. Neither opening (average 24.3 ± 5.2 ms) nor closing (average 10.9 ± 4.4 ms) are ATP dependent. Large errors (s.d.) are due to low number of spontaneous opening events under these conditions.

**Figure 3S4:**
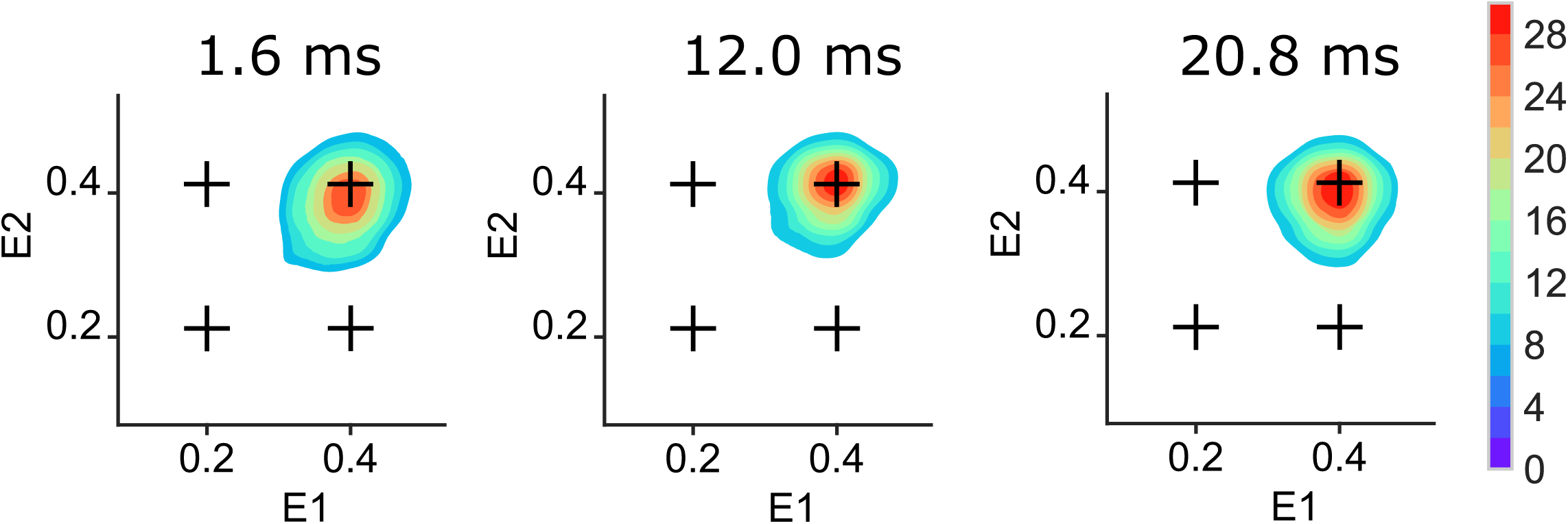
2D transition density plots in the presence of AMP-PNP. Time evolution of states for SecY_MK_EG:SecA:SecB:pOA in the presence of 1 mM AMP-PNP. The scale bar on the right depicts count contour levels.

**Figure 3S5:**
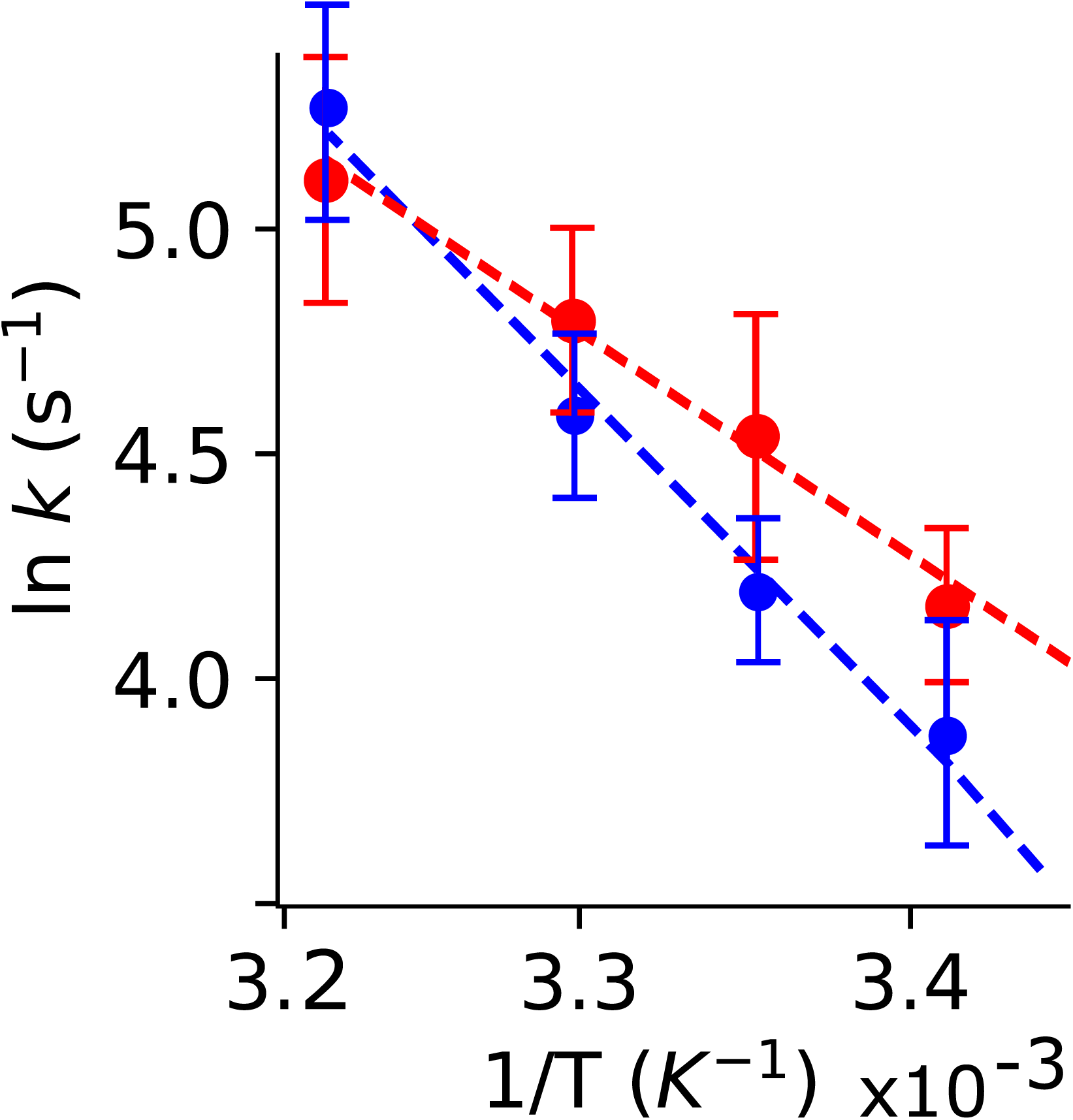
Activation energies for plug opening and closing. Arrhenius plot of reciprocal transition times (rates) for the plug opening (blue) and closing (red) under saturating 2 mM ATP concentration. Activation energy of 61.2 ± 4.5 kJ/mol for opening was obtained from the slope of a linear approximation (blue dashed line). The activation energy for plug closing is 45.1 ± 3.7 kJ/mol (red dashed line).

**Figure 5S1:**
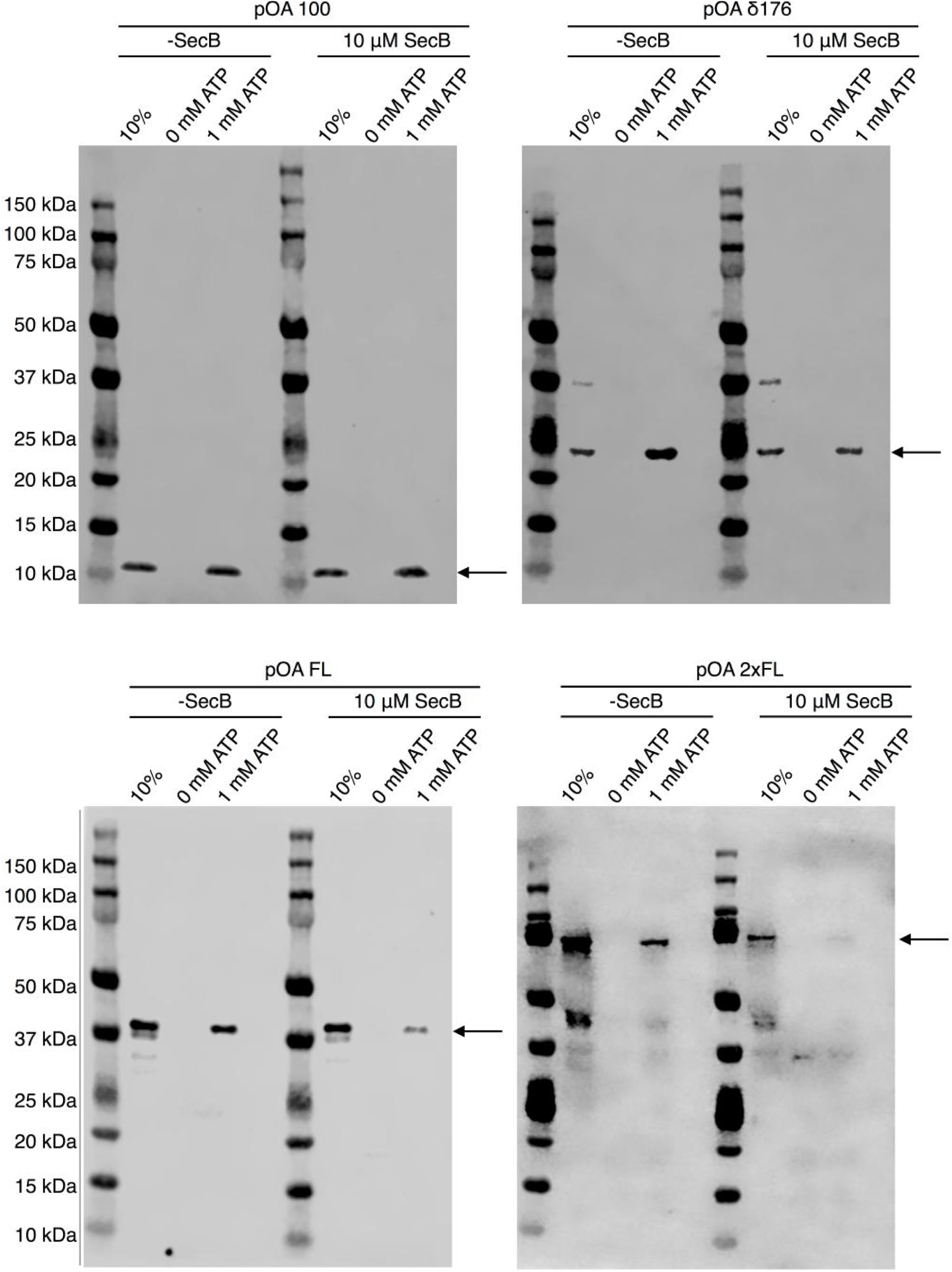
Translocation of pOA constructs with different lengths. Lengths are as follows: pOA (100 aa); pOA δ176 (233 aa); FL (354 aa); 2xFL (683 aa). Reactions were performed in the presence or absence of SecB. Gel lane: 10% of starting pOA loaded without protease treatment (positive and normalization control); control without ATP, translocation mix with ATP. Black arrows indicate the expected position of the translocated substrate.

**Figure 5S2:**
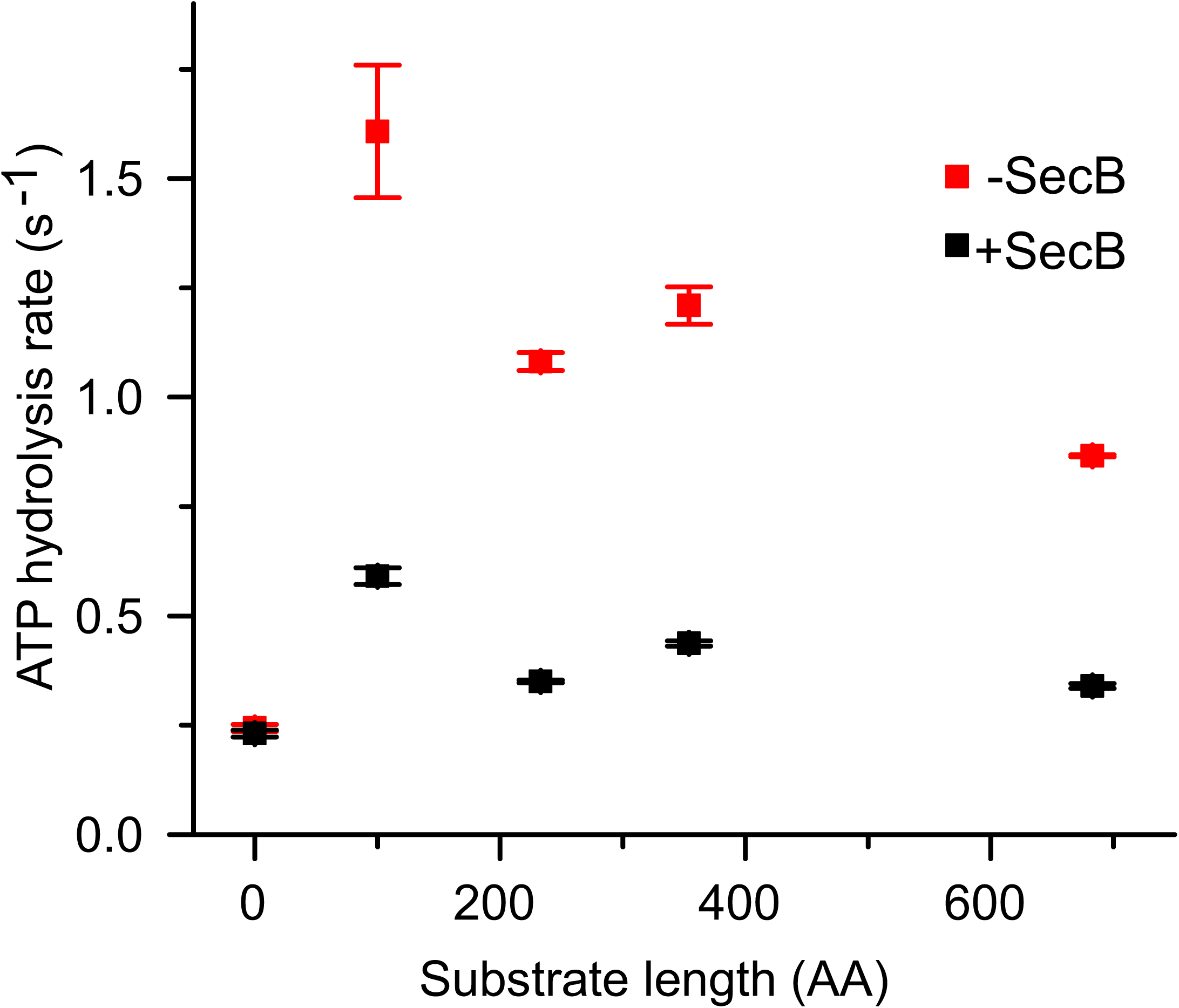
Ensemble ATPase activity stimulated by pOA constructs with different lengths. Ensemble ATPase activity stimulated by pOA constructs with different length in the presence (black) or absence (red) of SecB (10 µM). The rates were obtained under saturating ATP/pOA conditions. The zero-length substrate represents basal ATP hydrolysis activity by SecA alone.

**Figure 5S3:**
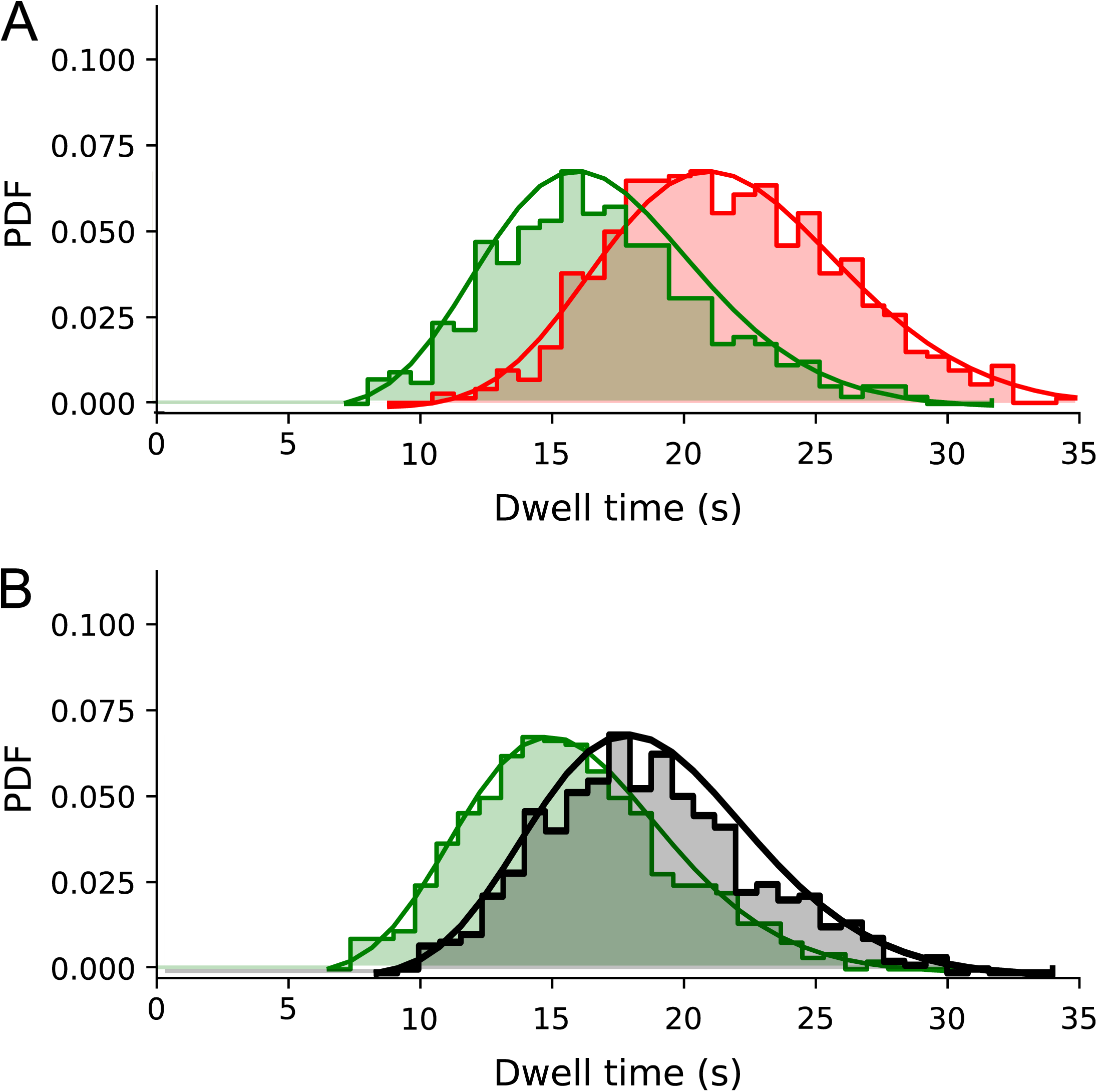
Deconvolution of photobleaching effect from the dwell time distributions for the longest 2xFL (683 aa) substrate. **A:** Without SecB, uncorrected distributions (shown as probability density functions) are shown in green, corrected distribution shown in red. Solid lines represent fitted gamma distribution functions. **B:** As A, but in the presence of SecB (10 µM). The corrected distribution is shown in grey. Solid lines represent fitted gamma distribution functions.

**Figure 5S4:**
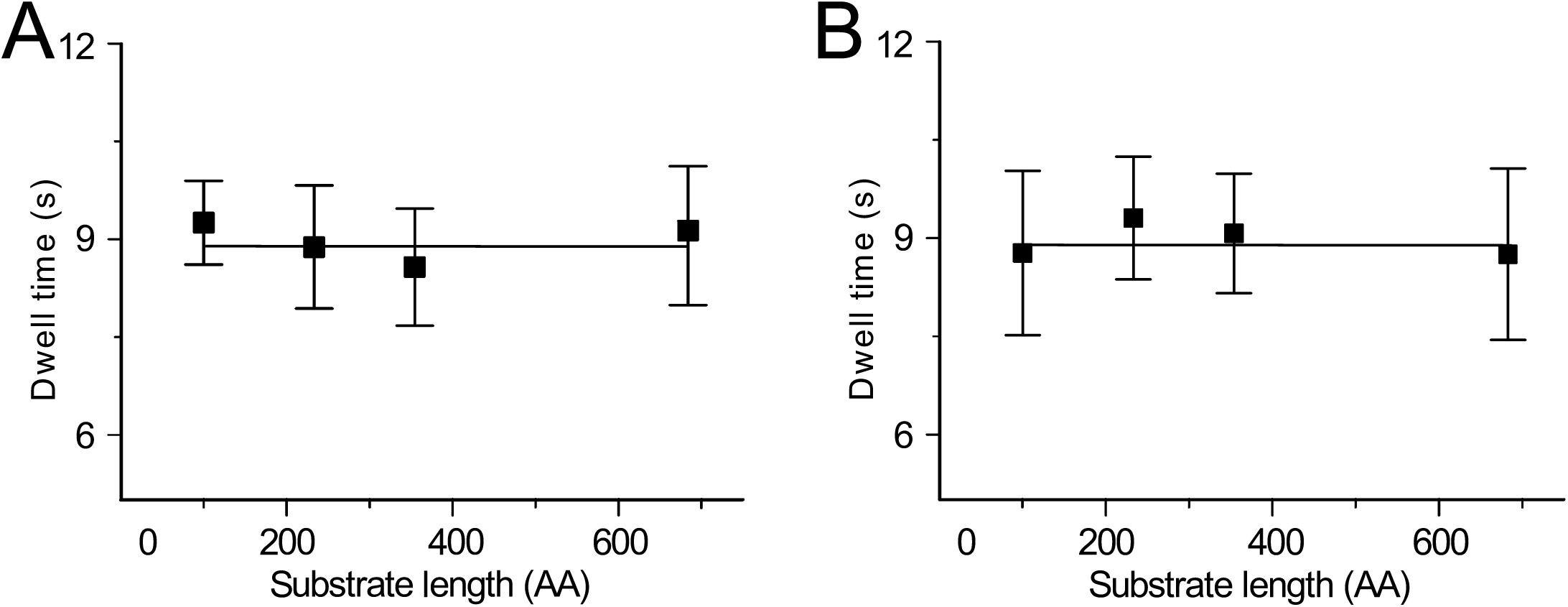
Dwell time of the closed *E*_*FRET*_ *~ 0.4* state as a function of the translocating substrate length. **A:** Without SecB. Error bars represent the standard deviation (s.d.) computed from the distribution of the dwell times. **B:** As **A**, but in the presence of SecB (10 µM).

